# Beyond the reference: gene expression variation and transcriptional response to RNAi in *C. elegans*

**DOI:** 10.1101/2023.03.24.533964

**Authors:** Avery Davis Bell, Han Ting Chou, Annalise B. Paaby

## Abstract

A universal feature of living systems is that natural variation in genotype underpins variation in phenotype. Yet, research in model organisms is often constrained to a single genetic background, the reference strain. Further, genomic studies that do evaluate wild strains typically rely on the reference strain genome for read alignment, leading to the possibility of biased inferences based on incomplete or inaccurate mapping; the extent of reference bias can be difficult to quantify. As an intermediary between genome and organismal traits, gene expression is well positioned to describe natural variability across genotypes generally and in the context of environmental responses, which can represent complex adaptive phenotypes. *C. elegans* sits at the forefront of investigation into small-RNA gene regulatory mechanisms, or RNA interference (RNAi), and wild strains exhibit natural variation in RNAi competency following environmental triggers. Here, we examine how genetic differences among five wild strains affect the *C. elegans* transcriptome in general and after inducing RNAi responses to two germline target genes. Approximately 34% of genes were differentially expressed across strains; 411 genes were not expressed at all in at least one strain despite robust expression in others, including 49 genes not expressed in reference strain N2. Despite the presence of hyper-diverse hotspots throughout the *C. elegans* genome, reference mapping bias was of limited concern: over 92% of variably expressed genes were robust to mapping issues. Overall, the transcriptional response to RNAi was strongly strain-specific and highly specific to the target gene, and the laboratory strain N2 was not representative of the other strains. Moreover, the transcriptional response to RNAi was not correlated with RNAi phenotypic penetrance; the two germline RNAi incompetent strains exhibited substantial differential gene expression following RNAi treatment, indicating an RNAi response despite failure to reduce expression of the target gene. We conclude that gene expression, both generally and in response to RNAi, differs across *C. elegans* strains such that choice of strain may meaningfully influence scientific conclusions. To provide a public, easily accessible resource for querying gene expression variation in this dataset, we introduce an interactive website at https://wildworm.biosci.gatech.edu/rnai/.

## Introduction

Research in the model organism *C. elegans* has yielded insight into myriad aspects of biology, particularly development, genetics, and molecular biology (Corsi et al., 2015). Historically, much of this work has been conducted in a single isogenic strain, the laboratory strain N2 (Andersen et al., 2012; Antoine Barriere & M. A. Felix, 2005). However, *C. elegans* harbors significant intraspecific genetic diversity (A. Barriere & M. A. Felix, 2005; Antoine Barriere & M. A. Felix, 2005; Crombie et al., 2019; Lee et al., 2021; Andersen et al., 2012), and in the last decade *C. elegans* has also been established as a powerful system for elucidating connections between genotype and phenotype (Andersen et al., 2012; Andersen & Rockman, 2022; A. Barriere & M. A. Felix, 2005; Antoine Barriere & M. A. Felix, 2005; Cook et al., 2017; Crombie et al., 2019; Evans, van Wijk, et al., 2021; Gaertner & Phillips, 2010; Lee et al., 2021). Natural genetic variation exists for practically any organismal trait measurable in *C. elegans* (Andersen & Rockman, 2022), for example: responsiveness to toxins, metals, drugs, and other stressors (Dilks et al., 2021; Evans & Andersen, 2020; Evans, Wit, et al., 2021; Hahnel et al., 2018; Na et al., 2020; Webster et al., 2019; Zdraljevic et al., 2019; Zdraljevic et al., 2017); behavior (Bendesky et al., 2012; Ghosh et al., 2015; McGrath et al., 2009); transgenerational mortality traits (Frezal et al., 2018; Saber et al., 2022); and efficiency in RNA interference (RNAi) (Elvin et al., 2011; Felix, 2008; Felix et al., 2011; Paaby et al., 2015; Tijsterman et al., 2002).

Naturally, molecular phenotypes that act as intermediaries between genotype and organismal traits, such as gene expression, also vary across strains. Studies from recombinant inbred lines (Evans & Andersen, 2020; Rockman et al., 2010; Vinuela et al., 2010) and, more recently, RNA sequencing of 207 wild strains (Zhang et al., 2022), have identified numerous expression quantitative trait loci (eQTL) that encode differences in gene expression. How such expression differences manifest across different strains, whether they offer clues into functional differentiation, and how genetic differences compare to environmentally induced differences in gene expression or mediate gene expression responses to environmental stimuli remain interesting questions. These questions require genome-wide characterization of gene expression in multiple strains under multiple conditions.

One phenomenon of particular interest is RNA interference, a mechanism of gene expression regulation triggered by environmental or endogenous sources of double stranded RNA with broad-reaching influence over diverse aspects of organismal biology (Billi et al., 2014; Wilson & Doudna, 2013). RNAi was discovered in *C. elegans* (Fire et al., 1998), but competency in response to environmental triggers is highly variable across wild *C. elegans* strains (Elvin et al., 2011; Felix, 2008; Felix et al., 2011; Paaby et al., 2015; Tijsterman et al., 2002). Previous work showed that a loss-of-function mutation in Argonaute RNAi effector gene *ppw-1* is largely responsible for the near-complete failure of Hawaiian strain CB4856 to mount an RNAi response against germline targets (Tijsterman et al., 2002), and later work characterized the failure in CB4856 as a much delayed, rather than absent, response (Chou et al., 2022). Other strains incompetent for germline RNAi exhibit distinct modes of RNAi failure with distinct genetic bases (Chou et al., 2022; Elvin et al., 2011; Pollard & Rockman, 2013). Even as wild strains vary in overall competency for germline RNAi, strain-to-strain differences in RNAi phenotypic penetrance are also highly dependent on the target gene; whether these differences arise from strain-specific developmental consequences of gene knock-down or strain-specific differences in target-dependent RNAi efficacy is unclear (Paaby et al., 2015). How this phenotypic variation in RNAi response is reflected in genome-wide transcriptional changes upon RNAi induction remains a largely open question.

Here, we evaluate how genotype (strain) and induction of the RNAi response affect the *C. elegans* transcriptome. We also consider how reliance on the reference genome, derived from the laboratory strain N2, might constrain estimates of gene expression in wild strains, and how a focus on N2 in studies of RNAi might limit inferences about RNAi biology within *C. elegans* generally. To investigate these questions, and to provide a public resource for interrogating transcriptional variation in this system, we performed RNA sequencing on five *C. elegans* strains with varying competency in germline RNAi, both in the control condition and under RNAi treatment targeting two germline-expressed genes.

## Materials and methods

### Sample preparation and sequencing

#### Worm strains and husbandry

Strains used in this study include wild strains CB4856, EG4348, JU1088, and QX1211 (gifts from Matthew Rockman) and wild-type laboratory strain N2 (gift from Patrick McGrath). Worms were cultured under standard conditions (Stiernagle, 2006) except that plates used for non-N2 wild strains were made with 1.25% agarose to prevent burrowing. All strains except for QX1211 were maintained at 20°C; QX1211 was maintained at 18°C to prevent induction of its mortal germline phenotype (Frezal et al., 2018). Worms were maintained for at least three generations without starvation before RNAi induction and RNA sequencing.

#### RNA interference

RNAi was induced via feeding and was carried out on plates at 20°C following established methods (Ahringer, 2006; Kamath et al., 2001). Worms were fed HT115 *E. coli* bacteria that had been transformed with the empty pL4440 vector or the pL4440-derived vectors *par-1* (H39E23.1) and *pos-1* (F52E1.1) from the Ahringer feeding library (Kamath & Ahringer, 2003). Bacteria cultures were prepared by streaking from frozen stocks onto LB agar with carbenicillin (25 ug/mL) and tetracycline (12.5 mg/mL); next 5-10 colonies from < 1 week old plates were used to inoculate liquid cultures of LB broth with carbenicillin (50 ug/mL) and tetracycline (12.5 mg/mL), which were then incubated with shaking at 37°C for 16-18 hours and finally amplified with carbenicillin (50 ug/mL) for 6hrs at a 1:200 dilution. 10cm agar feeding plates with 1mM IPTG (Ahringer 2006) were seeded with the RNAi bacteria cultures, then used within 44-78 hours after incubation in the dark. Worm strains reared under standard conditions were bleached on day 1 to synchronize, then bleached again on day 4 (Stiernagle, 2006). On day 5, L1s were transferred to the RNAi plates. All strains were exposed to RNAi in this way at the same time in triplicate, 6 total plates per strain.

#### RNA library preparation and sequencing

As previously described (Chou et al., 2022), synchronized hermaphrodites reared on RNAi feeding plates were washed off at the first sign of egg laying, washed twice with M9 buffer, and stored in TRIzol (Invitrogen #15596026) at -80°C until RNA extraction. RNA was extracted from all samples at the same time using TRIzol (Invitrogen #15596026) and RNeasy columns (Qiagen #74104) following (He, 2011). cDNA and sequencing libraries were generated from 500 ng of fresh RNA samples with 10 cycles of PCR with the NEBNext Ultra II Directional RNA Library Prep Kit for Illumina (NEB #7760). After quality checking using an Agilent 2100 Bioanalyzer, library fragments were size-selected via BluePippon (Sage Science). Single-end 75bp reads were sequenced on an Illumina NextSeq at the Molecular Evolution Core facility at the Georgia Institute of Technology.

### Analysis

#### Analytical approach

We considered multiple state-of-the-art pipelines to align RNA-seq data and quantify expression. Because the four wild strains in our study are diverged from the N2 reference genome by differing degrees (Cook et al., 2017), we required a method that could evaluate N2 data and non-N2 data over a range of variation without bias. One variant-aware option for quantifying RNA expression is to consider only RNA-seq reads that align to exactly one position on the reference genome (unique mappers) using STAR (Dobin et al., 2012), and to discard reads not uniquely aligning to the same position after non-reference variants are swapped into the read using WASP (van de Geijn et al., 2015). We explored this approach with our data. Specifically, we used STAR v2.7.5a with non-default parameters --outFilterMismatchNmax 33 – seedSearchStartLmax 33 --alignSJoverhangMin 8 --outFilterScoreMinOverLread 0.3 -- alignIntronMin 40 --alignIntronMax 2200 --waspOutputMode SAMtag --varVCFfile <VCF containing SNPs from all 4 non-reference strains>; these latter parameters implemented WASP from within STAR.

A second option is to generate strain-specific transcriptomes that incorporate known variants from each strain into the reference genome and use those to quantify transcript expression via pseudo-alignment; this approach permits reads to map to multiple locations (Bray et al., 2016; Patro et al., 2017). We do not compare the STAR-WASP approach to this pseudo-alignment approach here; high-level results were similar between the approaches. For our final analysis we chose the second option, for multiple reasons: 1) pseudo-alignment approaches are at least as accurate at estimating expression while being computationally more efficient (Bray et al., 2016; Patro et al., 2017); 2) pseudo-alignment approaches take into account the large fraction of reads that align to multiple loci in the genome (Bray et al., 2016; Patro et al., 2017); and 3) our specific generation of strain-specific transcriptomes enabled us to include insertion-deletion polymorphisms (INDELs), whereas WASP ignores INDELs (van de Geijn et al., 2015). Including INDELs was particularly relevant in this study, as 8,195-67,267 INDELs differentiate the four non-reference strains from the reference genome (CeNDR 20210121 release) (Cook et al., 2017).

The following methods detail generation of strain-specific transcriptomes and pseudo-alignment to quantify expression at individual genes. A subset of these methods and data overlap with our recent RNAi-focused study, which examined expression variation at specific RNAi genes (Chou et al., 2022).

#### Strain-specific transcriptomes

As previously described (Chou et al., 2022), we used SNPs and INDELs from CeNDR (release 20210121) (Cook et al., 2017) to update the N2 reference genome (release ws276) (Harris et al., 2020) to generate strain-specific transcriptomes using the software g2gtools (v0.1.31 via conda v4.7.12, Python v2.7.16) (https://github.com/churchill-lab/g2gtools). Specifically, INDELS were added to the reference genome with *g2gtools vcf2chain* and SNPs with *g2gtools patch*.

INDELs were added to the SNP-updated genome with *g2gtools transform*. We generated strain-specific GTFs from the strain-specific FASTAs with *g2gtools convert* and generated strain-specific transcriptomes from these GTFs with gffread (v0.12.7) (Pertea & Pertea, 2020).

The nextflow workflow performing this process is available in this project’s code repository (https://github.com/averydavisbell/wormstrainrnaiexpr) in *workflows/strainspectranscriptome*.

#### Gene expression quantification

Transcript-level quantification, used downstream for gene-level estimates, was performed using Salmon (v1.4.0) (Patro et al., 2017), as we previously detailed (Chou et al., 2022). First, we trimmed Illumina TruSeq adapters from RNA-seq reads with Trimmomatic (v0.3.9) (Bolger et al., 2014), parameters *ILLUMINACLIP:TruSeq3-SE.fa:1:30:1.* Strain-specific transcriptomes were used to generate Salmon index files with command *salmon index* with options *-k 31 -- keepDuplicates* (all others default; no decoy was used). Salmon transcript quantification *salmon quant* was performed with options *-l SR --dumpEq, --rangeFactorizationBins 4, --seqBias, and -- gcBias*, and library-specific fragment length arguments *--fldMean* and *--fldSD*.

The nextflow workflow generating strain-specific transcriptomes (link above) also generates strain-specific salmon indexes; the nextflow workflow performing transcript quantification is available in this project’s code repository in *workflows/strainspecsalmon*.

#### Differential expression analysis

Differential expression analyses were performed in R (v4.1.0) (R Core Team, 2021) using the DESeq2 package (v1.32.0) (Love et al., 2014). We imported transcript quantification data into DESeq2 using the tximport package (v1.20.0) (Soneson et al., 2015), which adds Salmon-specific transcript length normalizations to DESeq2’s sample-wise RNA quantification normalization and converts Salmon’s transcriptome quantification estimates to gene-level quantification estimates. Genes with fewer than 10 estimated reads across all samples (summed) were excluded from downstream analyses, retaining 18,589 genes. Principal components analysis was performed using the top 500 most variably expressed genes across all samples after DESeq2’s variance-stabilizing transformation (*vst* function), which was performed blind to experimental design.

We used DESeq2’s likelihood-ratio tests to determine whether genes were differentially expressed based on strain in the control condition and whether the interaction of strain and treatment was significant. For strain-wise significance, control sample counts were modeled with the negative binomial model

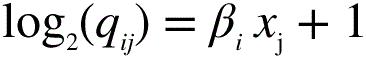

Which was compared to the reduced (null) model

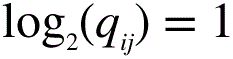

Here, for gene *i*, sample *j, q* is proportional to the actual concentration of RNA fragments for a gene (derived by DESeq2 from input counts and error modeling. (Love et al., 2014). *β_i_* gives the log2 fold changes for gene *i* corresponding to strain *x.* A total of 15,654 genes were sufficiently detected in the control samples to be included in this analysis (the remainder were excluded by DESeq2’s p-value correcting methods).

To evaluate strain:treatment interactions, all sample counts were modeled with the negative binomial model

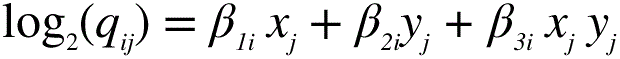

Which was compared to the reduced model

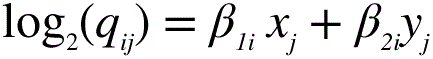

Here, the symbols are as in the first set of equations, with the additions that *y* corresponds to RNAi treatment; *xy* to the strain-treatment interaction; and *β_1_* to the strain effect, *β_2_* to the treatment effect, and *β_3_* to the interaction effect.

In both likelihood-ratio tests, genome-wide adjusted p-values were determined by DESeq2’s multiple testing correction. Genes were considered differentially expressed if this p-value was less than 0.1.

On the same datasets, we assessed differential expression within strains using DESeq2’s Wald’s tests of contrasts between treated (*par-1* or *pos-1* RNAi) and control (empty vector) samples. Genes were considered significantly differentially expressed if, after log2 fold change shrinkage using the ‘ashr’ method from the package ashr (v2.2-47) (Stephens, 2017), their absolute value fold change was greater than 1.5 and genome-wide adjusted p-value (FDR) was less than 0.1.

The script performing these analyses is available in this project’s code repository at *diffexp_lrt_straintreat_salmon_deseq2.R*.

*DNA sequence coverage estimation and identification of low-coverage and missing genes* We examined DNA sequence coverage within genes in CeNDR (Cook et al., 2017) BAM files (20210121 release); these files correspond to the same strains as in our study except in the case of EG4348, where CeNDR sequenced genetically identical strain EG4349. We note, of course, that the CeNDR DNA alignments were made directly to the N2 genome; we used the variants discovered therein to build our genotype-specific pseudo-transcriptomes. To get per-gene DNA sequence coverage, we first generated a file containing the non-overlapping, non-duplicated locations of all genes’ RNA generating sequences by determining the locations of all merged exons genome-wide using GTFTools (v0.8.5) (Li, 2018) (http://www.genemine.org/gtftools.php). Then, we determined the mean per-base coverage of each of these regions using mosdepth v0.3.2 (Pedersen & Quinlan, 2018) with default options with the exception of setting *--flag 1540,* which excludes unmapped reads, PCR duplicates, and QC failures. Finally, we computed the per-gene coverage as

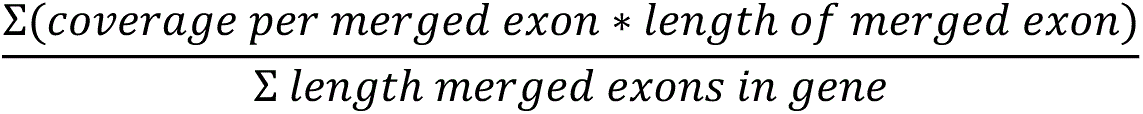

To delineate a set of low DNA coverage genes, we median-normalized the coverages within strain and flagged any with < 25% of the median coverage (i.e., median-normalized coverage < 0.25) as low coverage. Genes were classified as putatively missing from non-reference strain genomes if they had raw coverage estimates of exactly zero.

The workflow running this analysis is available in this project’s code repository in *workflows/mosdepthmergedexons*; this workflow performs custom gene-level analysis steps by calling an R script available in this project’s code repository at *exploregenecoverage_fromexons.R*. The scripts determining overlap with differentially expressed genes and zero-coverage genes are available in this project’s code repository at *de_dnacov_overlap.R* and *exploregencoverage_fromexons_lowend.R*.

#### ‘Off’ gene analysis

To identify genes putatively unexpressed in one or more strains despite being expressed in others (‘off’ genes), we first identified all genes differentially expressed between any two strains in the control condition (Wald’s test comparing each strain pair, genome-wide adjusted p < 0.1). The rationale was that genes significant for differential expression between strain pairs must have meaningful expression in at least one strain; we employed this standard to avoid inclusion of genes that are simply not expressed or expressed at a very low level regardless of strain. We then determined the average variance-stabilizing transformed (DESeq2 function *vst*) expression across all samples from all three treatments within each strain for these genes and identified those with zero mean expression. (These genes, of course, also have zero estimated expression prior to *vst* normalization.) Genes with strain-wise differential expression and zero expression within a strain comprise the ‘off’ gene set. (This process identified an additional six genes that fell just short of significance in the global analysis for differential expression in the likelihood-ratio test described above.) We then interrogated these genes for overlap with low DNA coverage and differential expression under RNAi treatment.

The script performing these analyses is available in this project’s code repository at *offgenes_straintreatDE_deseq2_dnacov.R*.

#### Gene set enrichment analysis

We performed gene set enrichment analysis of genes differentially expressed upon RNAi treatment using WormBase’s enrichment analysis tool (Angeles-Albores et al., 2016; Harris et al., 2020) (https://wormbase.org/tools/enrichment/tea/tea.cgi). We analyzed genes upregulated and downregulated on each RNAi treatment in all five strains (20 analyses total; 5 strains x 2 treatments x 2 directions of differential expression). Upregulated genes were those with higher expression on a treatment, with fold change > 1.5 vs control and adjusted p-value < 0.1; downregulated genes were those with lower expression on a treatment, with fold change < -1.5 vs control and adjusted p-value < 0.1 (see ‘Differential expression analysis’). The background gene set for all analyses was the 18,529 genes included in overall differential expression analyses. All gene-set enrichment related outputs were saved and the enrichment results tables (‘Download results table here’) output were combined across strains for visualization. The script performing this limited downstream processing is available in this project’s code repository at *exploreGeneSetEnrichmentResults.R*.

#### High-performance computation

Computationally intensive analyses were performed on the infrastructure of PACE (Partnership for an Advanced Computing Environment), the high-performance computing platform at the Georgia Institute of Technology. These analyses comprised pseudo-transcriptome generation, expression quantification, DNA sequence coverage estimation, and their related computational tasks.

#### Figures and website

Figures were made in R (v4.1.0) (R Core Team, 2021) using packages ggplot2 (v3.3.6) (Wickham, 2016), data.table (v1.14.3) (Dowle & Srinivasan, 2022) (https://r-datatable.com), DESeq2 (v1.32.0) (Love et al., 2014), cowplot (v1.1.1) (Wilke, 2020), ggVennDiagram (v1.2.0) (Gao, 2021), eulerr (v6.1.1) (Larsson, 2021), and ggpattern (v1.0.1) (FC et al., 2022), with color schemes developed using RColorBrewer (v1.1-3) (Neuwirth, 2022) and Paul Tol’s color palettes (https://personal.sron.nl/~pault/). The interactive website that enables exploration of the data from this study was developed using Shiny (Chang et al., 2022).

## Results and discussion

To investigate natural variation in both gene expression and response to exogenous RNAi, we performed RNA sequencing on five isogenic *C. elegans* strains in three conditions: RNAi targeting the germline genes *par-1* and *pos-1* and the untreated condition. We included the RNAi-competent reference strain N2 and four wild strains with varying competency to germline RNAi (Paaby et al 2015, Chou et al 2022): JU1088 (highly competent), EG4348 (moderately competent), and CB4856 and QX1211 (largely incompetent). These wild strains also vary in divergence from N2, representing some of the least (JU1088) and most (QX1211) divergent strains (variants per kilobase vs. N2 genome: 0.82, 1.40, 1.99, and 4.20, respectively, from *Caenorhabditis elegans* Natural Diversity Resource [CeNDR] data (Cook et al., 2017)). To limit bias arising from differences between non-N2 sequencing reads and the N2 reference genome in our analysis, we first created strain-specific transcriptomes by inserting known single nucleotide and insertion/deletion variants from CeNDR (Cook et al., 2017) into the reference genome. Then, we pseudo-aligned the RNA reads to these strain-specific transcriptomes to quantify per-gene RNA expression in each strain on each condition, and estimated differential expression based on strain, RNAi treatment, and their interaction.

### Genotype (strain)-wise expression variation predominates, nominates functionally diverged genes

Overall, genotypic differences between strains explained more gene expression variation than RNAi treatment. We detected nominal expression at 18,589 genes across the full dataset; a principal components analysis of the 500 most variable genes shows distinct strain-wise partitioning of the variation (**Figure 1A**). To identify genes with significant expression differences between strains in just the control condition, we compared a model with a term for strain to one without (via a likelihood-ratio test) for each gene. Of the 15,654 genes included in this control-specific analysis, 5355, or approximately 34%, were differentially expressed across the five strains (likelihood-ratio test, genome-wide adjusted *p* < 0.1) (**File S1**). This fraction of genes with expression differences between strains is consistent with recent findings that 28% of assayed genes were associated with mappable genetic differences (eQTLs) across 207 wild strains (Zhang et al., 2022). Other systems, such as flies, also harbor extensive variation in gene expression: a recent study of 200 inbred *Drosophila melanogaster* strains detected strain-wise expression variation at the majority of genes (Everett et al., 2020). The experimental and analytical approach matters a great deal; in the *Drosophila* study, many more variable genes were identified using RNA-seq data than microarray data, and only 30-40% of differentially expressed genes were associated with mappable eQTLs (Everett et al., 2020).

**Figure 1.**
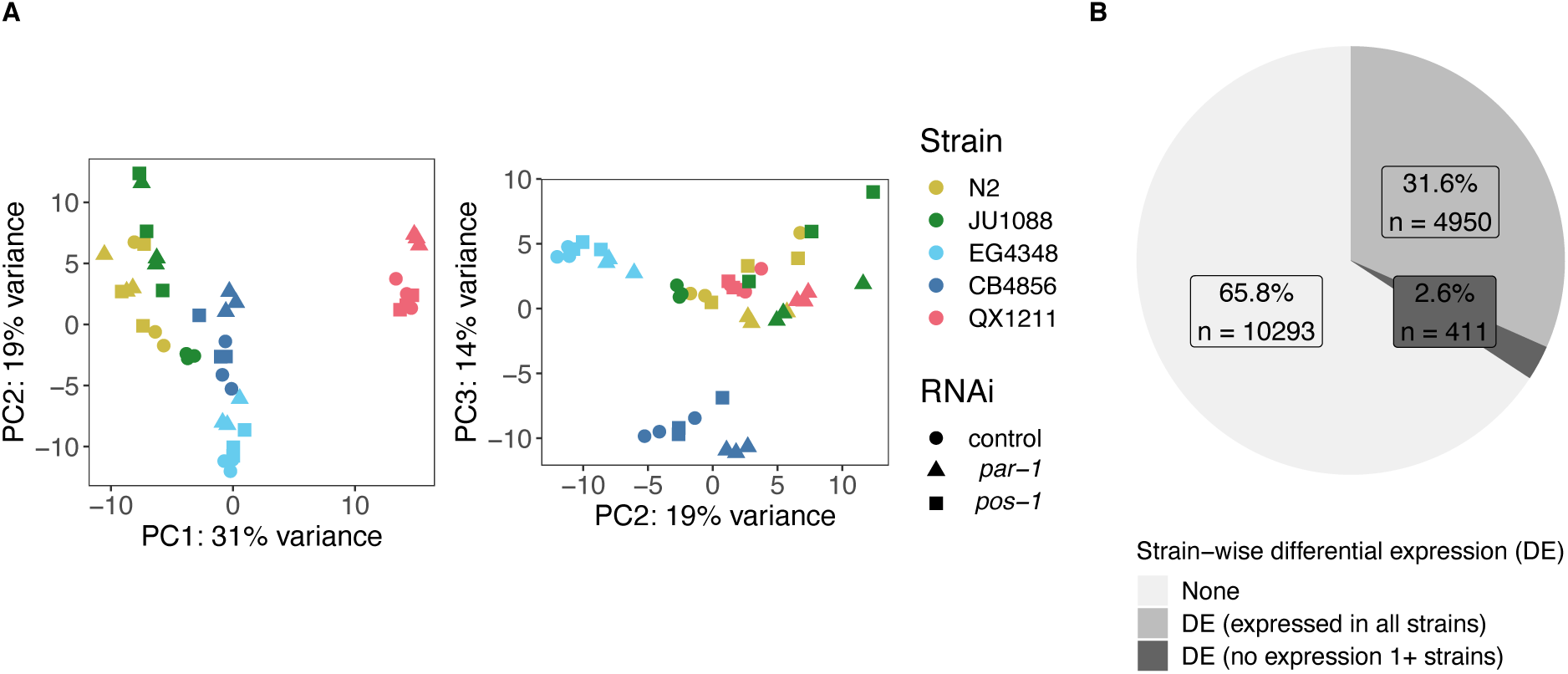
Genotype (strain) dominates expression variation across five *C. elegans* strains treated with RNAi targeting the genes *par-1* and *pos-1* or an empty vector control. **A)** Principal components analysis (PCA) of gene expression. PCs 1 vs. 2 (left) and 2 vs. 3 (right) of PCA of the 500 most variably expressed genes are plotted; the proportion of variance explained is noted on the axes. **B)** In the control condition, 34.2% of 15,654 nominally expressed genes are differentially expressed across strains (genome-wide adjusted *p* < 0.1 in a likelihood-ratio test between models including and excluding the *strain* term); a subset of these (approximately 2.6% overall) are not expressed at all in at least one strain (in any condition, see text for details). *Related Supplementary Material:* File S1 contains the genes differentially expressed based on strain File S2 contains the ‘off’ genes identified as potentially unexpressed in one strain but expressed in others

In some cases, presence versus absence of expression may underpin differential expression across strains; this pattern could indicate strain-wise differences in functional requirements or in developmental timing of expression. We identified such ‘off’ genes as those with zero mean expression in at least one strain (across all conditions) as well as significant strain-wise differential expression between a pair of strains in the control condition (genome-wide adjusted *p* < 0.1). This conservative zero-read threshold reduces the frequency of misclassifying low expression genes as off; the requirement for differential expression ensures true expression in at least one strain. This stringent selection yielded 411 putative ‘off’ genes (**Figure 1B**, **File S2**). Most of these genes lacked expression in a single strain: 249 were off in one strain, 105 were off in two strains, 51 were off in three strains, and only 6 genes were expressed in a single strain and off in the others (**Figure S1A**). We detected 49 genes that were off in N2 but expressed in at least one other *C. elegans* strain. The complete functional repertoire of these genes would therefore be invisible in a study using only the N2 strain. Such on/off patterns of gene expression occur in other systems as well; for example, across 144 *Arabidopsis thaliana* strains, thousands of genes showed strong expression in some strains but zero expression in others (Zan et al., 2016).

To assess the potential significance of ‘off’ genes in the context of RNAi response, we investigated whether any genes unexpressed in one strain exhibited differential expression within another strain following *par-1* or *pos-1* RNAi treatment. Of the 411 ‘off’ genes, 47 were differentially expressed on an RNAi treatment in at least one other strain (RNAi differential expression threshold: genome-wide adjusted *p* < 0.1 and fold change > 1.5 for within-strain RNAi treatment vs. control comparisons) (**Figure S1B**). The majority (*n* = 33) of these genes were differentially expressed in only one RNAi treatment in one strain. However, one gene identified by this analysis is W06G6.11 (WBGene00012313), which was ‘off’ in N2 but expressed in the other strains, and was significantly upregulated on RNAi against both *par-1* and *pos-1* in RNAi-sensitive strain JU1088 (fold change = 1.9 and genome-wide adjusted *p* = 0.03; fold change = 3.4 and genome-wide adjusted *p* = 0.003, respectively). Prior RNA-seq and microarray studies have indicated that W06G6.11 expression may be affected by the activity of Argonaute *alg-1* (Aalto et al., 2018), a member of the RNA-induced silencing complex involved in endogenous and exogenous short RNA processing (Grishok et al., 2001), and also by exposure to pathogens (Engelmann et al., 2011; Lee et al., 2013). These studies detect W06G6.11 expression in N2, but in samples derived from older adult hermaphrodites relative the young adults we sampled; a study that included CB4856 also confirmed significantly higher W06G6.11 expression in that strain relative to N2 (Zamanian et al., 2018).

This process of identifying genes that are unexpressed in some strains, but differentially expressed based on a treatment or phenotype of interest in others, might be used to identify candidate genes for other naturally variable phenotypes, perhaps as a complement to genotype-to-phenotype mapping by genome-wide association studies with expression mediation analyses (Evans & Andersen, 2020; Zhang et al., 2022).

### Reference bias screening increases confidence in differential expression calls

For RNA-seq studies that evaluate wild strains, reliance on a reference strain poses a concern. The main issue is whether the mapping of fewer non-reference strain RNA reads than reference-strain reads to a gene arise from true differences in gene expression, or from failure of non-reference reads to correctly map to the reference genome due to sequence divergence (reference bias) (Degner et al., 2009). Such discrepancies might remain even after the use of genotype-specific transcriptomes. In the case of *C. elegans*, wild strains exhibit a wide range in levels of divergence from the reference strain N2 in the species generally and the strains studied here specifically (Andersen et al., 2012; Cook et al., 2017; Crombie et al., 2019); much of this diversity is located in hyper-divergent haplotypes encompassing 20% of the genome (Lee et al., 2021).

To refine our level of confidence in the genes we identified as differentially expressed, we examined our results in the context of alignment quality in the original CeNDR genome sequencing data (Cook et al., 2017) (**Figure S2**, **Files S3, S4**). For each strain in our study, we curated a list of genes with missing or poor DNA sequence alignment in CeNDR (Cook et al., 2017) (**File S5**). Specifically, we classified genes with exactly zero coverage as missing in that strain’s genome; this is a conservative assignment, as even one well-aligned DNA sequence

Were differentially expressed genes associated with poor DNA coverage? Overall, yes: overlap of the missing-or-low coverage and strain-wise differentially expressed gene sets revealed significant enrichment (hypergeometric test of enrichment *p* = 9.8 x 10^-46^). However, the absolute number of differential expression genes with poor DNA coverage was modest: only 4% of all genes analyzed and 8% of genes with differential expression across strains had missing or low DNA coverage (**Figure 2B**). Put another way, 52% of missing or low DNA coverage genes were called as differentially expressed, while 29% of all analyzed genes were called as differentially expressed. Further, we note that poor DNA coverage arises from several sources. First, by chance, some genes will be low coverage simply due to stochastic variation in short-read sequencing depth, as reflected in the 62 genes binned as low coverage in N2 mapped to itself (**Figure 2A**). Second, sequence divergence between the mapped strain and the reference genome could inhibit alignment (reference bias); this possibility motivates this analysis. Third, the gene could be missing from the strain’s genome while present in the N2 reference genome. Not surprisingly, QX1211, the strain most diverged from the N2 reference genome, exhibits the most missing and the most low coverage genes (**Figure 2A**, **File S6**).

**Figure 2.**
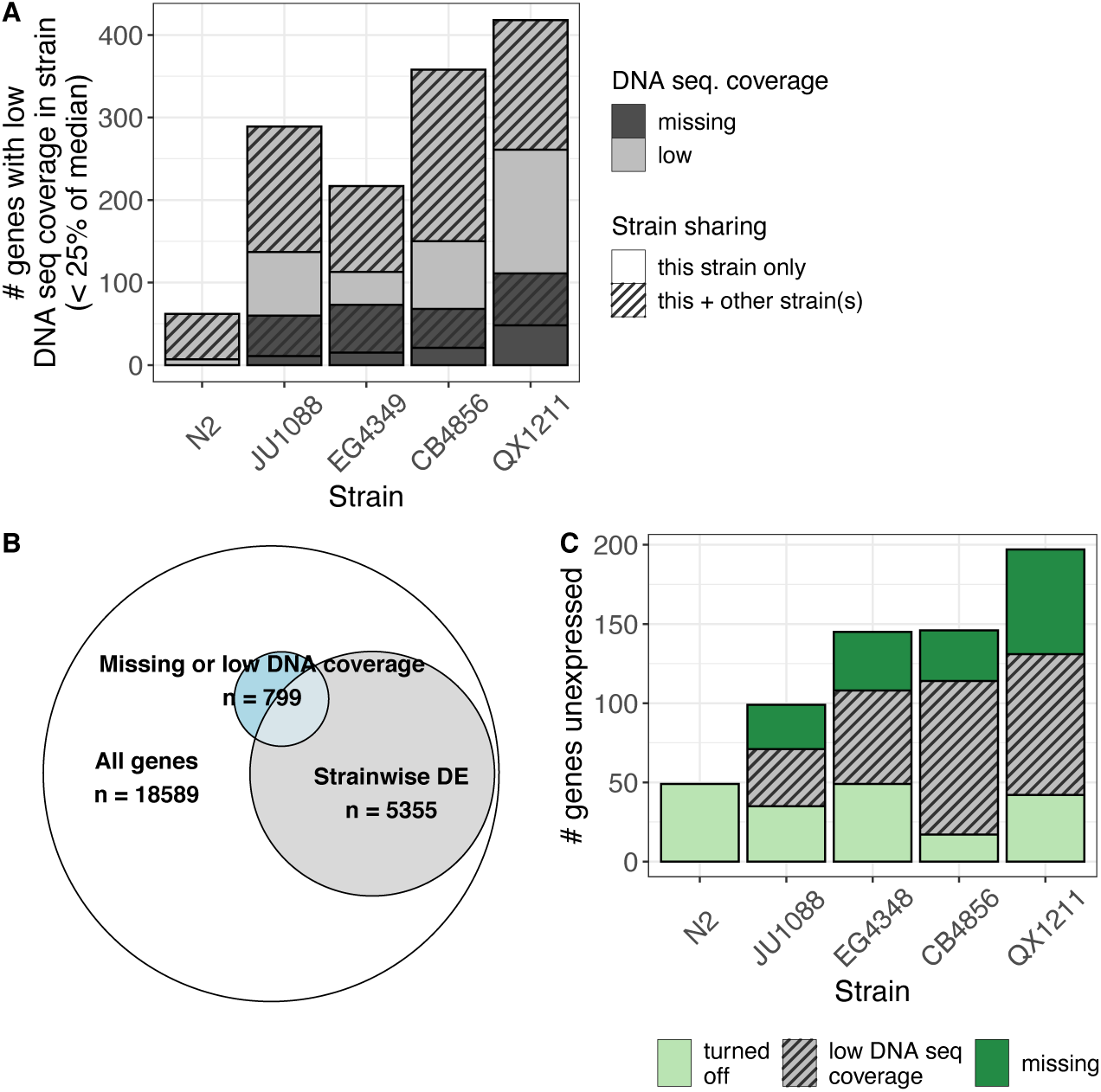
Improving confidence in differential expression calls by integrating DNA alignment data. **A)** The number of genes with low (<25% of the median) and missing (zero raw coverage) DNA alignment coverage (from CeNDR sequencing (Cook et al., 2017)) in each strain, of the 18,589 genes included in the expression analysis. Strain note: CeNDR assessed DNA coverage in EG4349, the genetically identical isotype to EG4348. **B**) The total number of genes differentially expressed based on strain (likelihood-ratio test of models including and excluding *strain* term, genome-wide adjusted *p* < 0.1) and their overlap with genes classified as missing or low DNA coverage in any strain (417 are both differentially expressed across strains and low DNA coverage, hypergeometric enrichment test *p* = 9.8 x 10^-46^). Areas are proportional to number of observations. **C)** The number of unexpressed ‘off’ genes per strain, subset into three categories: called as turned off at the RNA level with high confidence; missing in the strain genome (zero raw coverage); called with uncertainty, given low DNA sequence coverage (<25% but >0 median DNA coverage). *Related Supplementary Material:* Figure S2 shows DNA coverage distributions and cutoffs File S2 contains details on each ‘off’ gene File S3 contains raw per-gene DNA sequence coverage estimates File S4 contains median-normalized per-gene DNA sequence coverage estimates Files S5 contains the list of genes flagged as low DNA coverage *Files S6-7 provide numerical summaries of ‘off’ genes* read precluded a gene from being classified as missing. We classified genes with more than zero coverage but less than 25% of the gene-wise median DNA coverage in each strain as low coverage. This process identified a similar set of genes across strains despite the contribution of some strain-to-strain coverage variation (**Figure S2**, **File S5**). In total, we identified 799 genes as missing or low DNA coverage in one or more strains (**Figure 2A**).

The set of ‘off’ genes that show zero expression in some strains may be particularly vulnerable to reference bias, for example if they were more likely to be pseudogenes in at least one strain. In this scenario, poor DNA coverage may be conflated with true expression loss, as accumulated mutations may lead both to poor DNA coverage and consequently poor RNA alignment and to reduced expression through mutation-mediated defunctionalization. Here, when genes are detected as unexpressed, we can make distinctions between 1) missing genes, which we are reasonably confident do not exist in the strain genome; 2) genes for which we may not trust the conclusion of zero expression because of low DNA coverage and potential bias in RNA read mapping; and 3) true ‘off’ genes, which do not fall into either category and likely represent unbiased expression differences at the RNA level. In this scheme, among the four non-reference strains, 17-49 (12-35%) of the originally detected ‘off’ genes are likely truly turned off, 28-66 (22-34%) appear missing from the strain genome, and 36-89 (36-66%) are undetected for an unknown reason but have low DNA coverage and may be influenced by reference bias (**Figure 2C**, **File S7**).

As we would expect, all 49 ‘off’ genes in the reference strain N2 were classified as truly unexpressed; none were missing or low coverage (**Figure 2C**). Of these, 22 are listed as pseudogenes on WormBase (Harris et al., 2020), and may represent alleles that have been pseudogenized in the N2 lineage but remain functional in other strains. One such candidate is the Argonaute ZK218.8 (WBGene00013942), which is expressed in strains CB4856 and QX1211 and may reflect functional diversification in RNAi processes across the population (Chou et al., 2022). Of the 47 ‘off’ genes with *par-1* or *pos-1* RNAi effects in another strain, a large majority (*n* = 39, 83%) were missing in the genome or were associated with low DNA coverage (**Figure S3**). This majority represents a slight enrichment relative to the proportion of missing or low coverage genes within the complete set of ‘off’ genes (286/411 or 70%) (one-sided proportion test with continuity correction: χ^2^ = 3.05, df = 1, *p* = 0.04). Enrichment of genome divergence among RNAi-responsive ‘off’ genes supports the hypothesis that genes associated with RNAi are evolving rapidly in *C. elegans* (Chou et al., 2022). By adding the missing and low DNA coverage filters, we infer that, of genes with an RNAi effect in another strain, zero (in N2) to 12 (in QX1211) were missing from the strain’s genome and 1-6 genes per strain were present but truly unexpressed at the RNA level. These genes might be the most interesting candidates for downstream expression-based study. This set includes the putative RISC-associated gene W06G6.11 (WBGene00012313) discussed above.

An alternative approach to handling reference bias is to side-step it by excluding transcripts associated with known (Lee et al., 2021) hyper-divergent haplotypes (Zhang et al., 2022). However, because 1) some genes in hyper-divergent regions had good DNA alignment with low SNP density and others outside the regions had no DNA coverage, and 2) our study focuses exclusively on genic regions, we chose a gene-level, strictly coverage-based approach for bias screening. Still, a limitation of our approach (and most others) is that it cannot identify bias associated with elevated RNA levels in diverged or duplicated haplotypes relative to the N2 haplotype. Such bias could occur if reads in non-reference strains come from a gene poorly represented or missing in the reference, which are then spuriously assigned to an incorrect gene with a better match. This type of bias is difficult to define, quantify, and exclude.

Additionally, as for any arbitrary threshold, our cutoff of < 25% median coverage likely produces a mix of false positives and negatives, *i.e.*, genes with low DNA coverage but accurate RNA alignments and genes above the coverage cutoff that are nevertheless skewed by reference bias. While those interested in specific genes would therefore do well to interrogate them further, the DNA coverage approach provides a useful quality control filter for initial analyses of differential expression.

### Complex genotype and target specificity in transcriptional response to RNAi

Wild *C. elegans* strains vary in response to exogenous RNA interference. In particular, strains differ widely in competence for RNAi against germline targets delivered by feeding, as measured by phenotypic consequences following putative target knockdown (Elvin et al., 2011; Felix, 2008; Felix et al., 2011; Paaby et al., 2015; Tijsterman et al., 2002). To assess the transcriptional response to RNAi in worms with variable germline RNAi competencies, we fed worms dsRNA targeting the maternal-effect embryonic genes *par-1* and *pos-1* as well as the empty vector control. Both genes are expressed in the mature hermaphrodite germline and are essential for embryonic viability; in competent animals, RNAi by feeding results in dead embryos (Paaby et al., 2015; Sijen et al., 2001). Gene expression knockdown of the targets themselves confirmed the previously observed differences in RNAi competency (Chou et al., 2022; Paaby et al., 2015): under *pos-1* RNAi, *pos-1* expression levels dropped the most in JU1088, followed by N2 and then EG4348; strains CB4856 and QX1211 showed no drop in expression (**Figure S4A, C**). RNAi against *par-1*, which induces a less lethal response (Chou et al., 2022; Paaby et al., 2015), resulted in a similar though less strong pattern of *par-1* knockdown (**Figure S4B, D**). These results confirm that strains differ in RNAi response and that the response was target-gene-specific; this target specificity was also evident transcriptome-wide.

To assess how strains vary in overall transcriptional response to RNAi, we identified changes in gene expression across treatments (*par-1* RNAi, *pos-1* RNAi, and the negative control) that differed across the five strains. Specifically, for each gene in the dataset, we asked whether a model with or without a strain x treatment interaction term better explained the pattern of expression (see Methods). Genome-wide, 842 genes (5% of those assayed) varied in RNAi response across strains (*i.e.,* had significant strain:treatment interaction via likelihood-ratio test, genome-wide adjusted *p* < 0.1) (**File S8**). We also identified, within each strain, differences in expression following *par-1* and *pos-1* RNAi relative to the control. The number of genes differentially expressed under RNAi treatment (genome-wide adjusted p < 0.1, fold change > 1.5) varied substantially across strains and as well as between the two treatments (**Figure 3A, Figure S5**, **Files S9a-j**).

**Figure 3.**
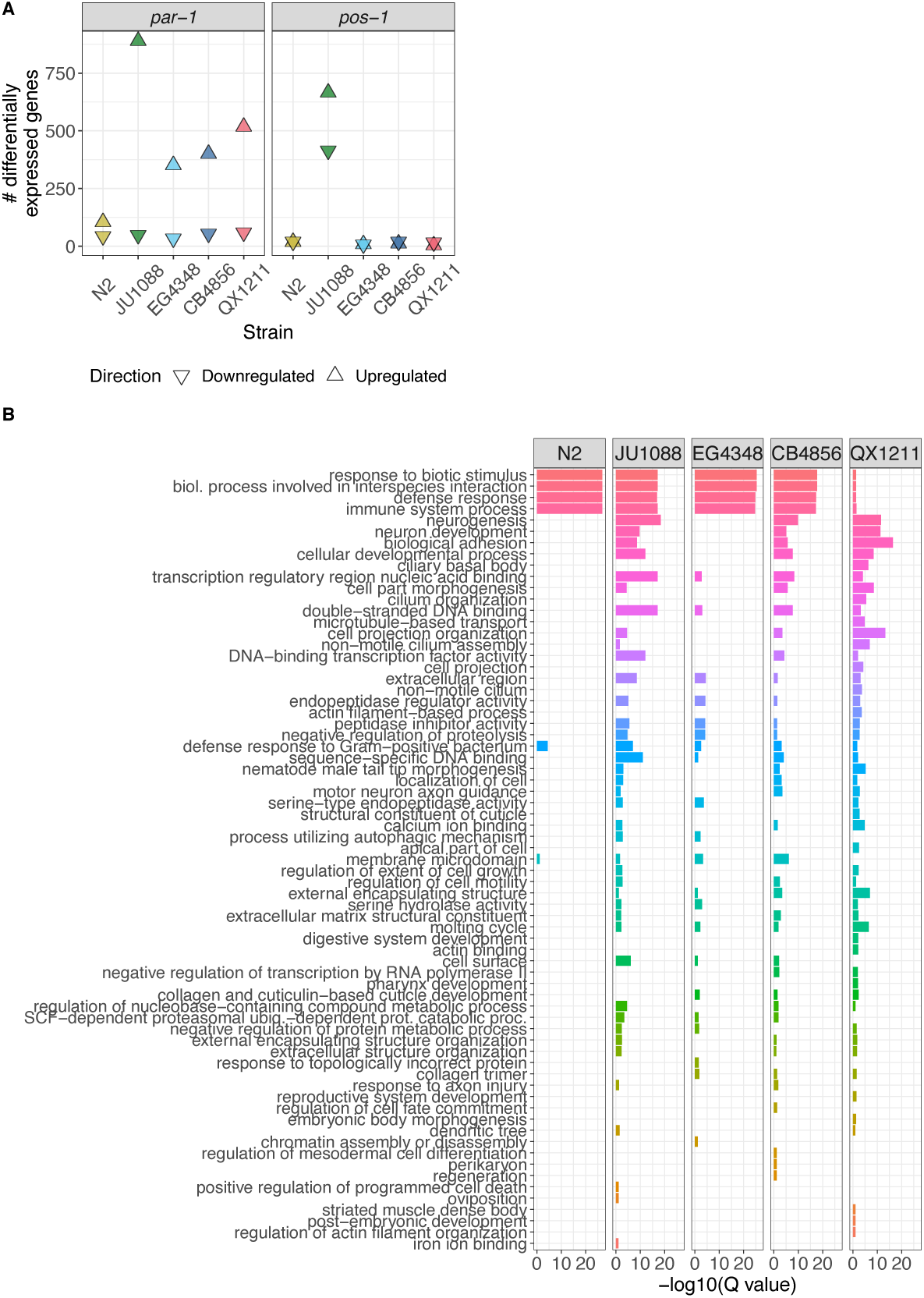
The transcriptional response to dsRNA is highly strain- and target-specific. **A)** The number of genes up- and down-regulated in each strain upon *par-1* and *pos-1* dsRNA ingestion/RNAi induction. Genes were called differentially expressed if their shrunken absolute fold change was > 1.5 and genome-wide adjusted p-value/FDR < 0.1. **B)** Gene set enrichment analysis results for genes upregulated on *par-1* dsRNA in each strain. Gene ontology (GO) categories that were significantly enriched (false discovery rate Q < 0.1) in any strain are included. GO terms are ranked and colored by median significance across strains. *Related Supplementary Material:* Figure S5 shows volcano plots for RNAi treatments for each strain Figure S6 contains Venn diagrams of overlap among strains in specific DE genes *Figure S7* *shows results from the same gene set enrichment analysis of genes downregulated under* par-1 *RNAi and up- and down-regulated under pos-1 RNAi* Table S1 gives number of up and downregulated genes in each strain and included in each analysis File S8 contains the genes differentially expressed based on strain-treatment interaction Files S9a-j contain the genes differentially expressed in each strain in each RNAi treatment vs. control File S10 gives all enriched GO categories.

On both *par-1* and *pos-1* RNAi, the highly germline-RNAi competent strain JU1088 exhibited the most differentially expressed genes relative to the control, suggesting that this strain is the most transcriptionally responsive to RNAi (**Figure 3A, Figure S5**). However, on *par-1* RNAi, the moderately competent strain EG4348 and the largely incompetent strains CB4856 and QX1211 showed substantially more differentially expressed genes than the competent laboratory strain N2. These results indicate that the number of genes transcriptionally responsive to exogenous RNAi is not predictive of RNAi phenotypic penetrance, and that ‘competence’ defined by end-point phenotypes and/or artificial triggers may obscure intermediary RNAi activity, or activity in alternative RNAi pathways (Chou et al., 2022).

Relative to *par-1*, *pos-1* RNAi induced substantially fewer differentially expressed genes in all strains but JU1088, indicating that RNAi transcriptional response is highly target-specific.

Furthermore, differential expression following *par-1* RNAi was strongly skewed towards an overabundance of upregulated genes compared to downregulated genes (**Figure 3A, Figure S5**). Of course, a transcriptional response may reflect developmental consequences of losing *par-1* or *pos-1* gene expression, at least in competent strains (Chou et al., 2022; Paaby et al., 2015); here, we cannot easily distinguish these effects from those arising from induction of the RNAi process itself. However, several lines of evidence suggest that RNAi process effects dominate. First, RNAi is a systemic phenomenon with a repertoire of many genes (Billi et al., 2014) while *par-1* and *pos-1* expression is largely restricted to the germline with consequential effects predominantly in the early embryo (Harris et al., 2020); our samples were prepared from whole worms. Second, the incompetent strains exhibited transcriptional responses genome-wide, but not at the targeted genes. Finally, as described below, the transcriptional response at a gene-by-gene level was strain-specific, consistent with our growing understanding of natural variation in RNAi.

To identify transcriptional responses to RNAi that may be universal within *C. elegans*, we first checked for differentially expressed genes that were shared across strains. However, overlap among strains was sparse (**Figure S6**): no genes with differential expression to both *par-1* and *pos-1* RNAi were shared across all five strains, and the only gene responsive to both treatments in the competent strains (JU1088, N2, and EG4348) was *asp-14*, a predicted aspartyl protease involved in innate immunity (Harris et al., 2020). Such strain-specific patterns fit with our observations of RNAi variability: not only does *C. elegans* exhibit substantial natural variation in germline RNAi competence (Elvin et al., 2011; Felix, 2008; Felix et al., 2011; Paaby et al., 2015;

Tijsterman et al., 2002), but the genetic basis for RNAi failure appears strain-specific as well (Chou et al., 2022). We posit that even among competent strains, *C. elegans* varies in details of the RNAi biological response mechanism, including which genes are affected, the magnitude or functionality of their activity, and their timing. These differences are apparent in the transcriptional responses of N2 and JU1088 (**Figure 3, Figure S6, Figure S7**), including the activity of W06G6.11 described above. As the RNAi response is also highly target-specific, these results portray RNAi as a phenomenon of exquisite specificity and context dependence.

However, statistical flux around significance cutoffs within strains may limit detection of gene-specific responses, and we also wished to examine the biological significance of the transcriptional responses. Therefore, we investigated whether the same general classes of genes responded to RNAi across strains by applying WormBase gene set enrichment analyses (Angeles-Albores et al., 2016; Harris et al., 2020) to the sets of genes up- and down-regulated on the RNAi treatments (**Files S9**). Strains showed a clear pattern of enriched gene ontology (GO) categories, particularly in the largest gene set, those upregulated under *par-1* RNAi (**Figure 3B**, **File S10**). Specifically, GO terms associated with canonical RNAi functions such as immune defense were well represented in all strains except in the germline incompetent strain QX1211, and genes in other categories were enriched in all strains except in N2. This pattern explains the paucity of differentially expressed genes in N2 relative to other strains following *par-1* RNAi (**Figure 3A**), as those in N2 are restricted to immunity associated ontology. These results demonstrate that reference strain N2 may not be a good representative for RNAi transcriptional response in *C. elegans* generally. Some of these patterns were also evident at genes downregulated under *par-1* RNAi, and up- and down-regulated under *pos-1* RNAi, though these results were less clear (**Figure S7**); this difference from *par-1* upregulated genes might reflect the more limited pool of differentially expressed genes in those categories.

In sum, transcriptional responses to RNAi differed across strains, but these responses did not clearly discriminate between RNAi competent and incompetent strains in the context of N2- derived GO categories: some competent strains upregulated non-defense categories while N2 did not, and incompetent strain CB4856 upregulated defense categories while incompetent strain QX1211 did not. That said, some strain-specific aspects of RNAi responses at the phenotype level may shed light on the transcriptional response enrichments. EG4348 is partially sensitive to RNAi (Chou et al., 2022; Felix et al., 2011; Paaby et al., 2015), and its GO term profile is similar to highly sensitive strain JU1088. While largely incompetent for germline RNAi, CB4856 does eventually exhibit strong RNAi phenotypes at late ages (Chou et al., 2022; Felix et al., 2011; Paaby et al., 2015; Tijsterman et al., 2002); its GO term profile similarity to JU1088 could be explained by the fact that this delay arises from the perturbation of a single gene, *ppw-1* (Tijsterman et al., 2002). Alternatively, QX1211 exhibits an apparent on/off response pattern among individual animals (Chou et al., 2022), and this binary penetrance of may be insufficient to detect defense/immune gene upregulation in a bulk analysis.

### A public web resource for data exploration

We have built a user-friendly, interactive website (https://wildworm.biosci.gatech.edu/rnai/) to enable straightforward public exploration of our gene expression data across the five wild *C. elegans* strains and three RNAi conditions. For any gene in our analysis, this website 1) visualizes the RNA quantification per sample split by treatment or strain, 2) allows the user to look up differential expression results between any two strain-treatment groups, 3) reports if expression differs by strain in the control condition and by RNAi treatment across strains, and 4) enables initial reference bias screening by displaying DNA sequencing coverage and whether the gene overlaps a hyperdivergent haplotype. This website may be useful for exploratory analyses of genes of interest for many types of studies in the *C. elegans* community.

## Conclusion

The results of the investigations described here further expand our understanding of *C. elegans* processes beyond the reference strain N2. Our quantification of gene expression variation among wild strains demonstrates that mapping bias arising from the use of a reference genome, while a greater liability for inferences about individual genes, can be restricted to a relatively minor concern for genome-wide studies in this system. However, the strain-specific variation in RNAi transcriptomic response suggests that our understanding of RNAi processes, derived predominantly from studies in N2, incompletely represents RNAi biology in *C. elegans* as a whole. The type of dataset presented here, genome-wide expression in multiple natural genetic backgrounds over multiple conditions of interest, enables researchers to characterize how much variation exists in the experimental systems we study. Understanding the scope of natural variation informs evolutionary hypotheses about traits of interest and offers insight into otherwise inaccessible relationships among genes, their functions, and phenotypes.

## Data availability

Strains and feeding vectors are available from CeNDR or the CGC, and upon request. All supplementary data files are available via Zenodo at https://doi.org/10.5281/zenodo.7406794: File S1 contains the genes differentially expressed based on strain; File S2 contains the ‘off’ genes identified as potentially unexpressed in one strain but expressed in others; File S3 contains raw per-gene DNA sequence coverage estimates; File S4 contains median-normalized per-gene DNA sequence coverage estimates; File S5 contains the list of genes flagged as low DNA coverage; Files S6-7 contain summaries of missing/zero coverage genes; File S8 contains the genes differentially expressed based on strain-treatment interaction; Files S9a-j contain the genes differentially expressed in each strain in each RNAi treatment vs. control; File S10 contains the results of the gene set enrichment analyses. Per-gene differential testing results and related information are available via an interactive web app at https://wildworm.biosci.gatech.edu/rnai/. Gene expression data (raw and processed) are available at GEO with the accession number GSE19083. Code used for all analyses can be found at https://github.com/averydavisbell/wormstrainrnaiexpr.

## Acknowledgments

We are grateful to members of the Paaby lab for helpful conversations about this work. We appreciate the use of the shared equipment, services, and expertise of the core facilities at the Parker H. Petit Institute for Bioengineering and Bioscience at the Georgia Institute of Technology. Specifically, we thank Shweta Biliya at the Molecular Evolution Core for collaboration on sequencing. Troy Hilley provided expert web server configuration support for the interactive web app. This research was supported in part through research cyberinfrastructure resources and services provided by the Partnership for an Advanced Computing Environment (PACE) at the Georgia Institute of Technology, Atlanta, Georgia, USA. As ever, WormBase served as an invaluable resource to this study. This research was funded by NIH grant R35 GM119744 to A.B.P. and NSF Postdoctoral Research Fellowship in Biology 2109666 to A.D.B.

## Supplementary Figures

**Figure S1.**
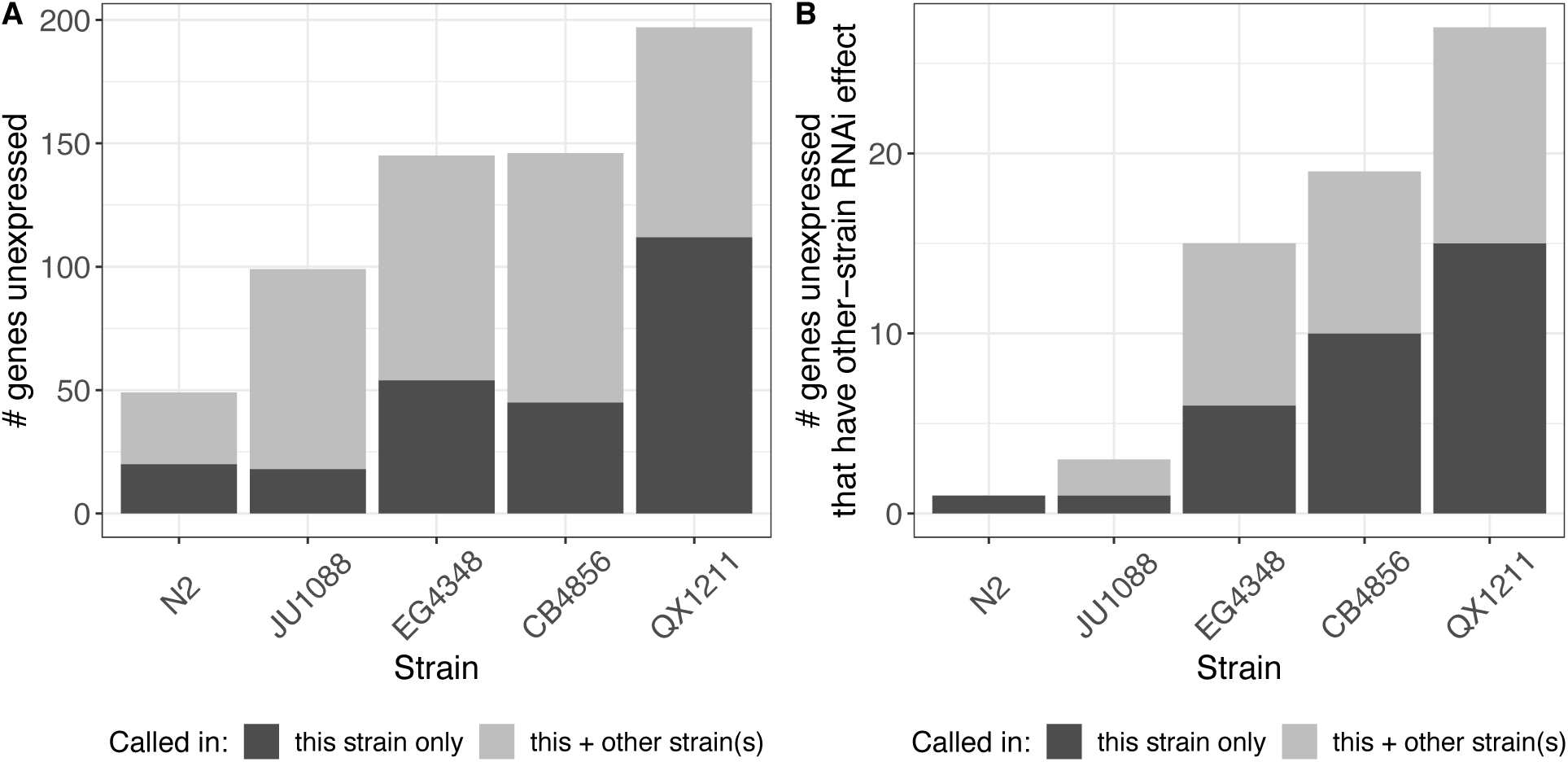
‘Off’ genes, which are expressed in at least one strain but show no expression in one or more others. **A)** All ‘off’ genes per strain, either unique or shared across strains (n = 411 total; genes may be present for multiple strains). **B)** The subset of ‘off’ genes that exhibit differential expression on RNAi to *par-1* or *pos-1* in other strains, which are potential candidates for RNAi functional divergence (n = 47 total; genes may be present for multiple strains). *File S2 contains identity and details for each of these ‘off’ genes*.

**Figure S2.**
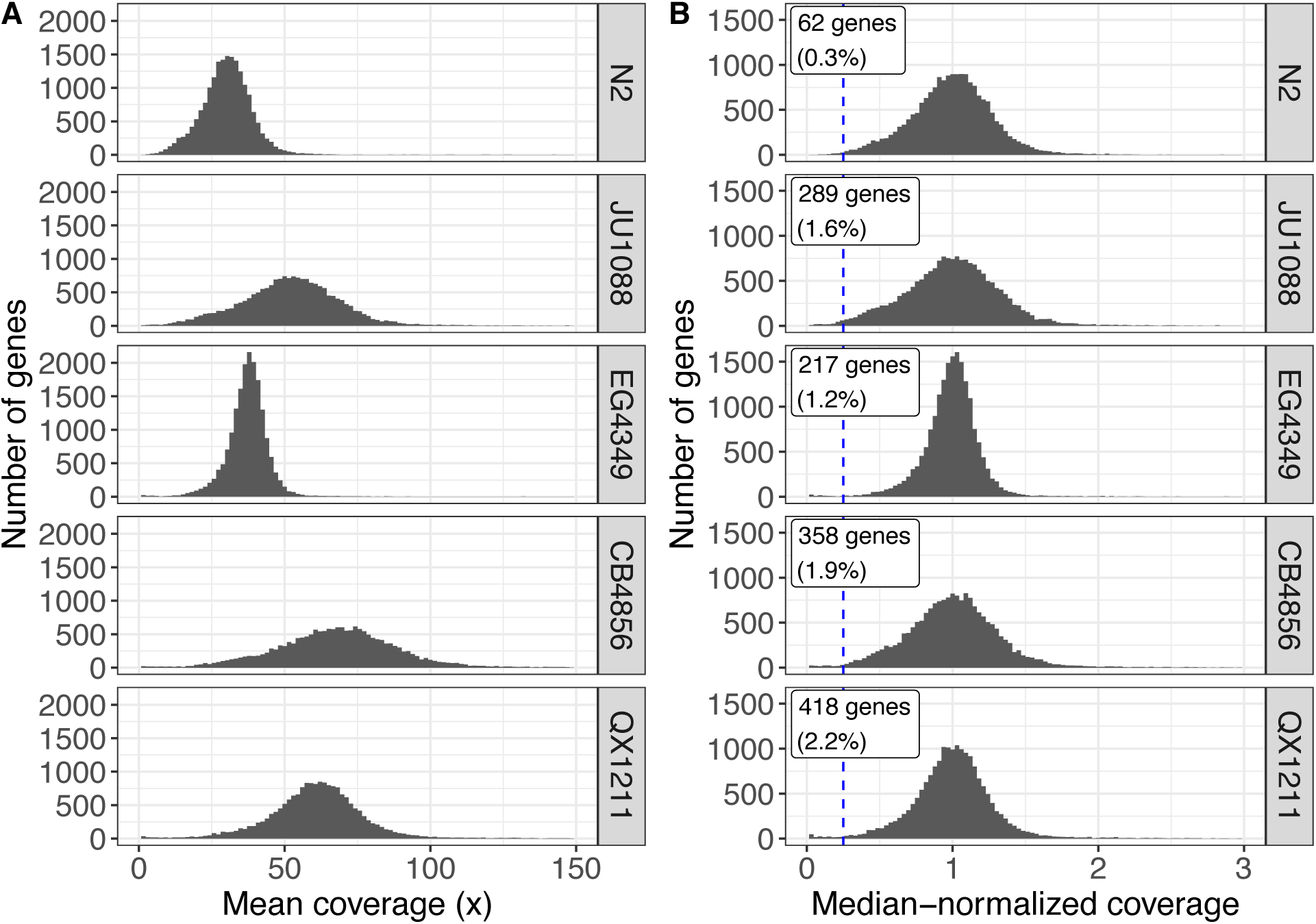
DNA sequence coverage across 18,589 genes included in expression analyses. Aligned DNA sequence data was obtained from CeNDR (release 20210121) (Cook et al., 2017). **A)** Mean coverage (mean number of reads covering each base) over merged non-overlapping exonic regions of genes in the five strains in this study. CeNDR assessed DNA coverage in EG4349, the genetically identical isotype to EG4348. The x-axis is truncated at 150x coverage for visual clarity, excluding 179 genes across all strains combined. **B)** Median-normalized coverage for the same genes as in **(A).** Genes with less than 25% median coverage are considered low DNA coverage in this study; this boundary is demarcated with the blue dashed line and the number and proportion of genes this set comprises is noted on the plots. The x-axis is truncated at 3x median coverage for visual clarity, excluding 227 genes across all strains combined. *Files S3 and S4 contain the source data. File S5 provides the list of genes identified as low coverage*.

**Figure S3.**
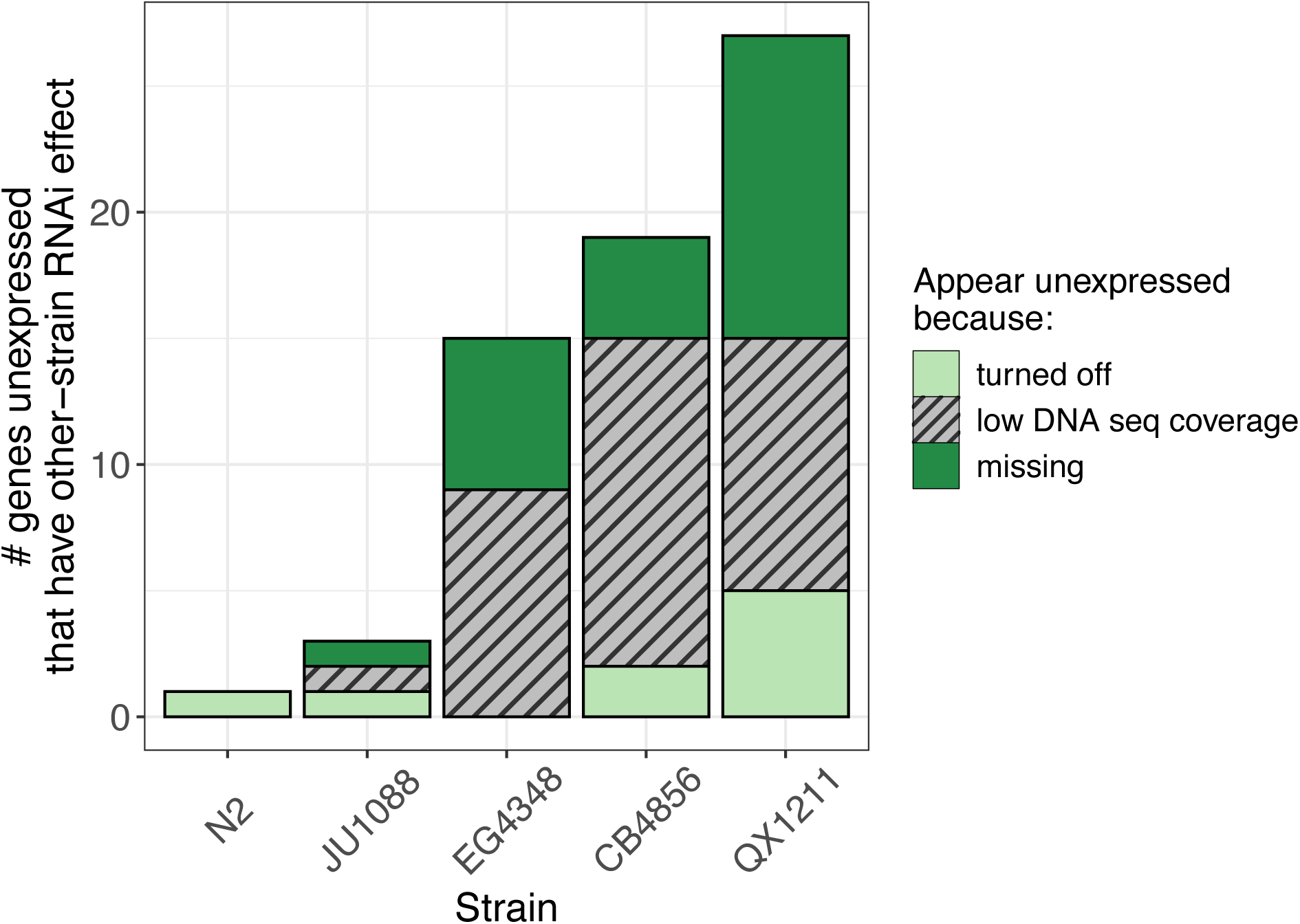
‘Off’ genes that were unexpressed in one or more strains but differentially expressed with respect to *par-1* or *pos-1* RNAi in another strain, potential candidates for RNAi functional divergence. DNA sequence coverage information is denoted with color and shading. Missing genes were those with zero DNA sequence coverage; low DNA sequence coverage genes had greater than zero but less than 25% median gene’s coverage; genes classified as truly turned off had greater than 25% median gene’s DNA sequence coverage. (DNA coverage was assessed in strain EG4349, isotype to EG4348).

**Figure S4.**
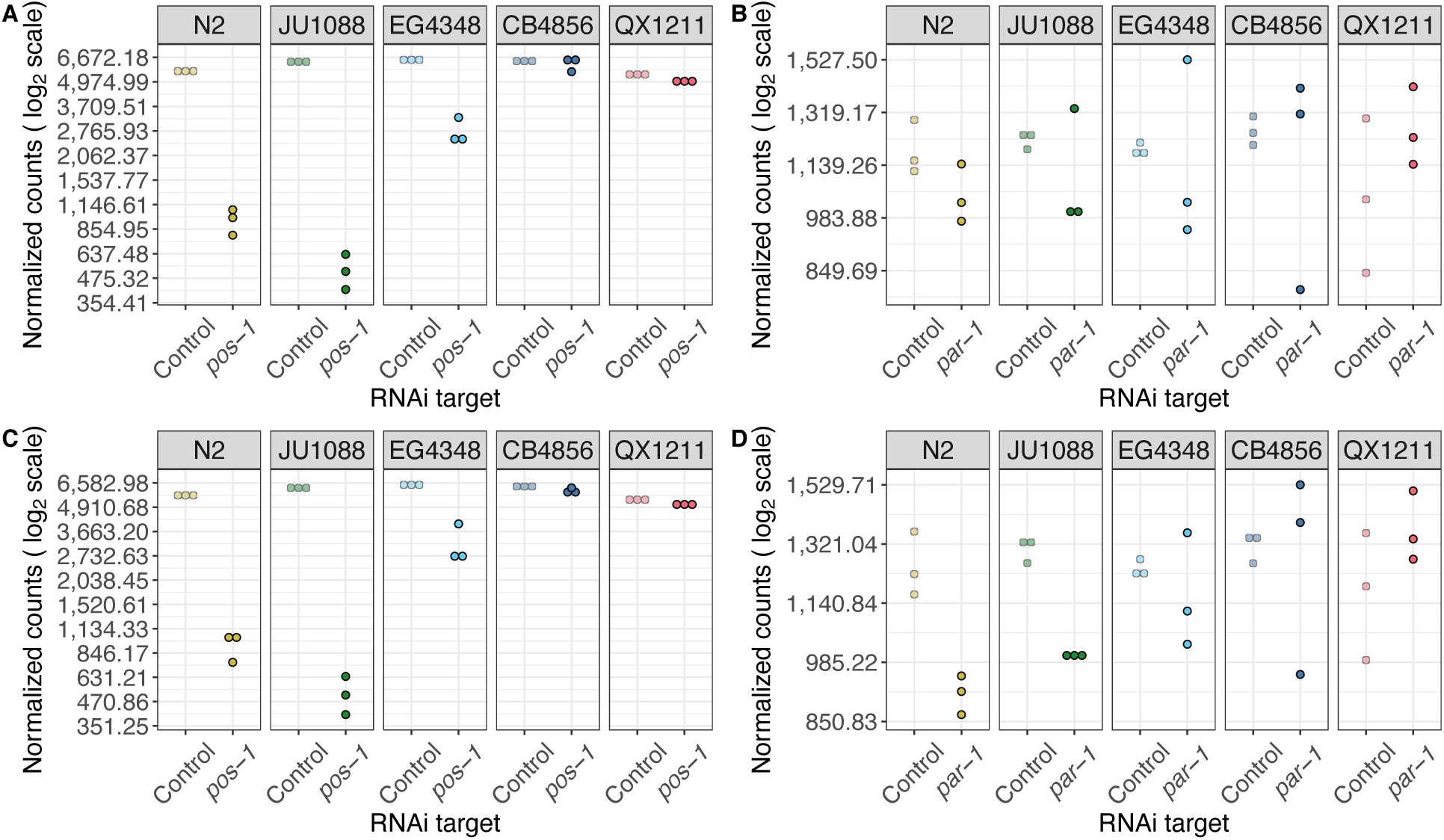
RNA-seq estimates suggest RNAi targets are knocked down commensurate with each strain’s RNAi capacity. **(A and B)** Quantification estimates from pseudoalignment to strain-specific transcriptomes, normalized to library size and gene length, as used for all analyses in this study. **A)** Quantification estimates for *pos-1* in control and exposure to *pos-1* dsRNA; response is significantly different across strains (the strain:treatment interaction is significant, genome-wide adjusted p = 4 x 10^-254^). **B)** Quantification estimates for *par-1* in control and exposure to *par-1* dsRNA (the strain:treatment interaction is not significant, genome-wide adjusted p = 0.92). **(C and D)** Detection of target knockdown is not dependent on RNAi strategy: panels show *pos-1* and *par-1* quantification estimates as in (**A and B**), respectively, but with alternative expression estimates derived from RNA sequence data uniquely mapping to one genomic location when containing the reference or non-reference allele (see methods).

**Figure S5.**
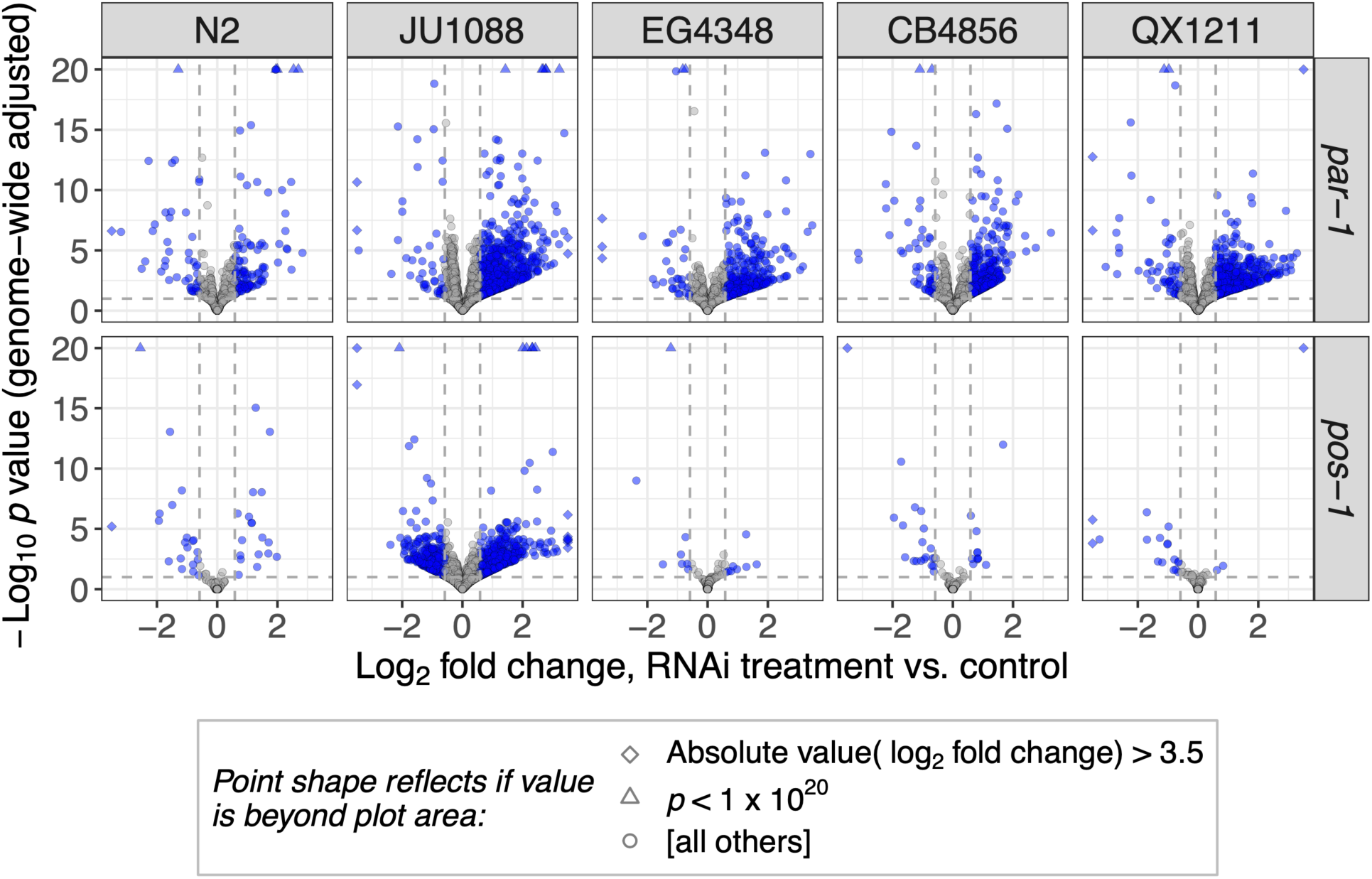
Volcano plots show genome-wide effects of RNAi treatments (against *par-1*, top, and *pos-1*, bottom) in each of the five strains. All genes with differential expression estimates are plotted; blue points denote genes with significant differential expression (genome-wide adjusted *p* < 0.1 and corrected [see methods] absolute value(fold change) > 1.5; these thresholds are annotated on the plot with gray dashed lines). For visual clarity, the y-axis is truncated at *p* = 10^-20^ and the x-axis is truncated at absolute log2 fold change = 3.5; genes with values exceeding these thresholds are included on the plots and are represented by unique point shapes as noted in the plot legend.

**Figure S6.**
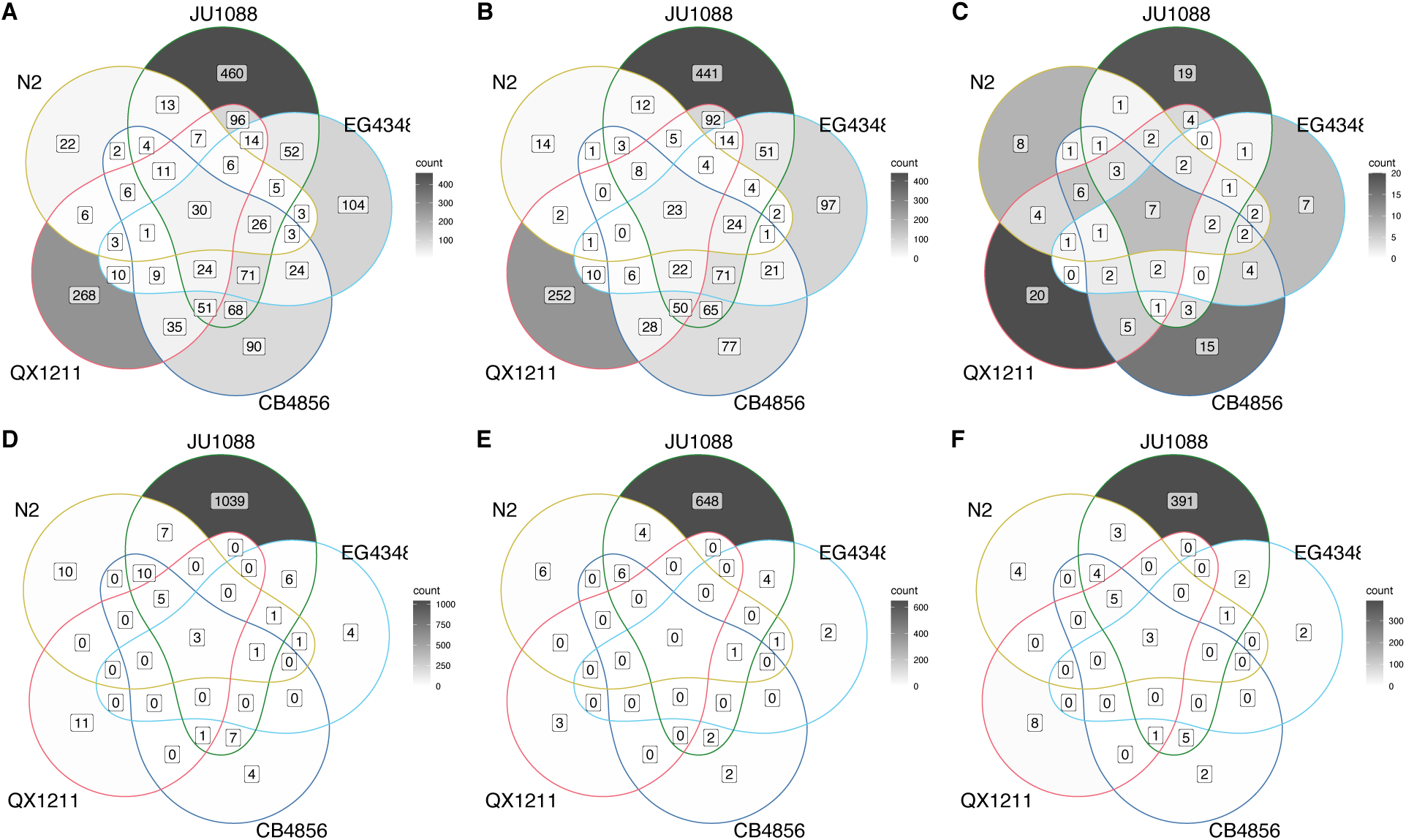
Limited overlap of genes called as differentially expressed in RNAi conditions vs. control across strains; shading scales with number of genes separately within each panel (see color bar legends). (**A-C)** Under *par-1* RNAi, genes differentially expressed in either direction **(A)**, upregulated **(B),** or downregulated **(C)**. **(D-F)** Under *pos-1* RNAi, genes differentially expressed in either direction **(D)**, upregulated **(E),** or downregulated **(F)**. Genes were called differentially expressed and included if their shrunken absolute fold change was > 1.5 and genome-wide adjusted p-value/FDR < 0.1 between RNAi and control within-strain. *Files S9a-j contain gene IDs and details.* Figure 3A *and Table S1 show the overall number of up- and down-regulated genes in each strain*.

**Figure S7.**
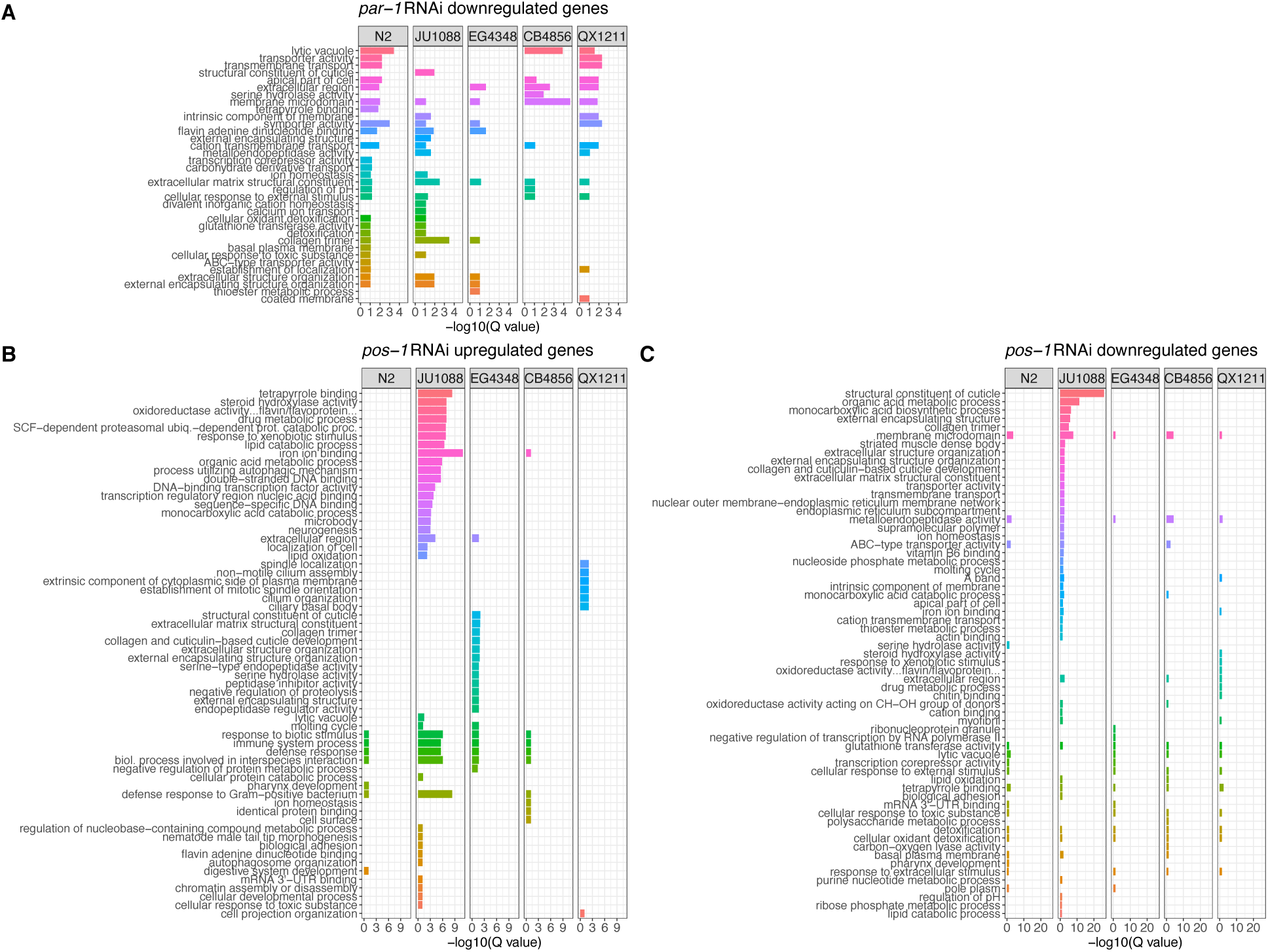
Gene set enrichment analysis results for genes **(A)** downregulated on *par-1* dsRNA in each strain, (**B**) upregulated on *pos-1* dsRNA, and (**C**) downregulated on *par-1* dsRNA. Only gene ontology (GO) categories significantly enriched (FDR Q < 0.1) in upregulated genes in any strain are included. GO terms are ranked and colored by median significance across strains. *Table S1 provides the number of genes included for each analysis. File S10 gives all enriched GO categories. Main* Fig 3B *displays the same analysis of genes upregulated under* par-1 *RNAi*.

## Supplementary Tables

**Table S1.**
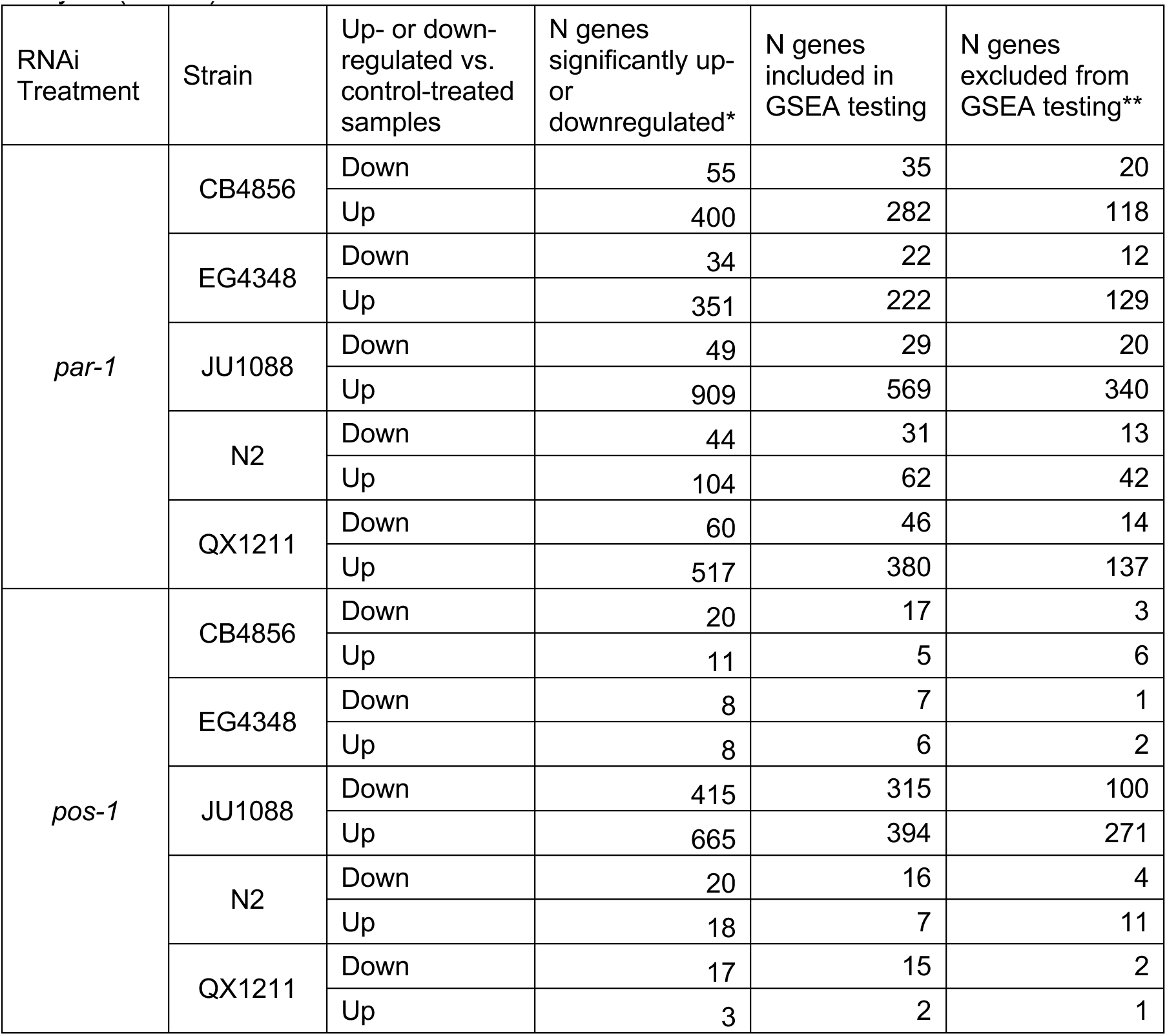
The number of genes differentially expressed in each RNAi treatment in each strain, relative to the control condition, as well as the number included in the gene set enrichment analysis (GSEA).

## Literature Cited

Aalto, A. P., Nicastro, I. A., Broughton, J. P., Chipman, L. B., Schreiner, W. P., Chen, J. S., & Pasquinelli, A. E. (2018, Jun). Opposing roles of microRNA Argonautes during Caenorhabditis elegans aging. PLoS Genet, 14(6), e1007379. https://doi.org/10.1371/journal.pgen.1007379

Ahringer, J. (2006). Reverse genetics. In V. Ambros (Ed.), WormBook. https://doi.org/doi/10.1895/wormbook.1.47.1

Andersen, E. C., Gerke, J. P., Shapiro, J. A., Crissman, J. R., Ghosh, R., Bloom, J. S., Felix, M. A., & Kruglyak, L. (2012, Jan 29). Chromosome-scale selective sweeps shape Caenorhabditis elegans genomic diversity. Nat Genet, 44(3), 285–290. https://doi.org/10.1038/ng.1050

Andersen, E. C., & Rockman, M. V. (2022, Jan 4). Natural genetic variation as a tool for discovery in Caenorhabditis nematodes. Genetics, 220(1). https://doi.org/10.1093/genetics/iyab156

Angeles-Albores, D., RY, N. L., Chan, J., & Sternberg, P. W. (2016, Sep 13). Tissue enrichment analysis for C. elegans genomics. BMC Bioinformatics, 17(1), 366. https://doi.org/10.1186/s12859-016-1229-9

Barriere, A., & Felix, M. A. (2005, Jul 12). High local genetic diversity and low outcrossing rate in Caenorhabditis elegans natural populations. Curr Biol, 15(13), 1176–1184. https://doi.org/10.1016/j.cub.2005.06.022

Barriere, A., & Felix, M. A. (2005). Natural variation and population genetics of Caenorhabditis elegans. WormBook. https://doi.org/10.1895/wormbook.1.43.1

Bendesky, A., Pitts, J., Rockman, M. V., Chen, W. C., Tan, M. W., Kruglyak, L., & Bargmann, C. I. (2012). Long-range regulatory polymorphisms affecting a GABA receptor constitute a quantitative trait locus (QTL) for social behavior in Caenorhabditis elegans. PLoS Genet, 8(12), e1003157. https://doi.org/10.1371/journal.pgen.1003157

Billi, A. C., Fischer, S. E. J., & Kim, J. K. (2014). Endogenous RNAi pathways in C. elegans. WormBook. https://doi.org/10.1895/wormbook.1.170.1

Bolger, A. M., Lohse, M., & Usadel, B. (2014, Aug 1). Trimmomatic: a flexible trimmer for Illumina sequence data. Bioinformatics, 30(15), 2114–2120. https://doi.org/10.1093/bioinformatics/btu170

Chang, W., Cheng, J., Allaire, J., Sievert, C., Schloerke, B., Xie, Y., Allen, J., McPherson, J., Dipert, A., & Borges, B. (2022). shiny: Web Application Framework for R. https://CRAN.R-project.org/package=shiny

Chou, H. T., Valencia, F., Alexander, J. C., Bell, A. D., Deb, D., Pollard, D. A., & Paaby, A. B. (2022). Diversification of small RNA pathways underlies germline RNAi incompetence in wild C. elegans strains. bioRxiv, 2021.2008.2021.457212. https://doi.org/10.1101/2021.08.21.457212

Cook, D. E., Zdraljevic, S., Roberts, J. P., & Andersen, E. C. (2017, Jan 4). CeNDR, the Caenorhabditis elegans natural diversity resource. Nucleic Acids Res, 45(D1), D650–D657. https://doi.org/10.1093/nar/gkw893

Corsi, A. K., Wightman, B., & Chalfie, M. (2015). A Transparent window into biology: A primer on Caenorhabditis elegans. WormBook. https://doi.org/doi/10.1895/wormbook.1.177.1

Crombie, T. A., Zdraljevic, S., Cook, D. E., Tanny, R. E., Brady, S. C., Wang, Y., Evans, K. S., Hahnel, S., Lee, D., Rodriguez, B. C., Zhang, G., van der Zwagg, J., Kiontke, K., & Andersen, E. C. (2019, Dec 3). Deep sampling of Hawaiian Caenorhabditis elegans reveals high genetic diversity and admixture with global populations. Elife, 8. https://doi.org/10.7554/eLife.50465

Degner, J. F., Marioni, J. C., Pai, A. A., Pickrell, J. K., Nkadori, E., Gilad, Y., & Pritchard, J. K. (2009, Dec 15). Effect of read-mapping biases on detecting allele-specific expression from RNA-sequencing data. Bioinformatics, 25(24), 3207–3212. https://doi.org/10.1093/bioinformatics/btp579

Dilks, C. M., Koury, E. J., Buchanan, C. M., & Andersen, E. C. (2021, Dec). Newly identified parasitic nematode beta-tubulin alleles confer resistance to benzimidazoles. Int J Parasitol Drugs Drug Resist, 17, 168–175. https://doi.org/10.1016/j.ijpddr.2021.09.006

Dowle, M., & Srinivasan, A. (2022). data.table: Extension of ‘data.fram’. https://Rdatatable.gitlab.io/data.table

Elvin, M., Snoek, L. B., Frejno, M., Klemstein, U., Kammenga, J. E., & Poulin, G. B. (2011, Oct 17). A fitness assay for comparing RNAi effects across multiple C. elegans genotypes. BMC Genomics, 12, 510. https://doi.org/10.1186/1471-2164-12-510

Engelmann, I., Griffon, A., Tichit, L., Montanana-Sanchis, F., Wang, G., Reinke, V., Waterston, R. H., Hillier, L. W., & Ewbank, J. J. (2011). A comprehensive analysis of gene expression changes provoked by bacterial and fungal infection in C. elegans. PLOS ONE, 6(5), e19055. https://doi.org/10.1371/journal.pone.0019055

Evans, K. S., & Andersen, E. C. (2020). The Gene scb-1 Underlies Variation in Caenorhabditis elegans Chemotherapeutic Responses. G3: Genes|Genomes|Genetics, 10(7), 2353-2364. https://doi.org/10.1534/g3.120.401310

Evans, K. S., van Wijk, M. H., McGrath, P. T., Andersen, E. C., & Sterken, M. G. (2021, Oct). From QTL to gene: C. elegans facilitates discoveries of the genetic mechanisms underlying natural variation. Trends Genet, 37(10), 933–947. https://doi.org/10.1016/j.tig.2021.06.005

Evans, K. S., Wit, J., Stevens, L., Hahnel, S. R., Rodriguez, B., Park, G., Zamanian, M., Brady, S. C., Chao, E., Introcaso, K., Tanny, R. E., & Andersen, E. C. (2021, Mar). Two novel loci underlie natural differences in Caenorhabditis elegans abamectin responses. PLoS Pathog, 17(3), e1009297. https://doi.org/10.1371/journal.ppat.1009297

Everett, L. J., Huang, W., Zhou, S., Carbone, M. A., Lyman, R. F., Arya, G. H., Geisz, M. S., Ma, J., Morgante, F., St Armour, G., Turlapati, L., Anholt, R. R. H., & Mackay, T. F. C. (2020, Mar). Gene expression networks in the Drosophila Genetic Reference Panel. Genome Res, 30(3), 485–496. https://doi.org/10.1101/gr.257592.119

FC, M., Davis, T. L., & ggplot2 authors. (2022). ggpattern: ‘ggplot2’ Pattern Geoms. https://CRAN.R-project.org/package=ggpattern

Felix, M. A. (2008, Nov 13). RNA interference in nematodes and the chance that favored Sydney Brenner. J Biol, 7(9), 34. https://doi.org/10.1186/jbiol97

Felix, M. A., Ashe, A., Piffaretti, J., Wu, G., Nuez, I., Belicard, T., Jiang, Y., Zhao, G., Franz, C. J., Goldstein, L. D., Sanroman, M., Miska, E. A., & Wang, D. (2011, Jan 25). Natural and experimental infection of Caenorhabditis nematodes by novel viruses related to nodaviruses. PLoS Biol, 9(1), e1000586. https://doi.org/10.1371/journal.pbio.1000586

Fire, A., Xu, S., Montgomery, M. K., Kostas, S. A., Driver, S. E., & Mello, C. C. (1998, Feb 19). Potent and specific genetic interference by double-stranded RNA in Caenorhabditis elegans. Nature, 391(6669), 806–811. https://doi.org/10.1038/35888

Frezal, L., Demoinet, E., Braendle, C., Miska, E., & Felix, M. A. (2018, Aug 20). Natural Genetic Variation in a Multigenerational Phenotype in C. elegans. Curr Biol, 28(16), 2588–2596 e2588. https://doi.org/10.1016/j.cub.2018.05.091

Gaertner, B. E., & Phillips, P. C. (2010, Dec). Caenorhabditis elegans as a platform for molecular quantitative genetics and the systems biology of natural variation. Genet Res (Camb), 92(5-6), 331–348. https://doi.org/10.1017/S0016672310000601

Gao, C.-H. (2021). ggVennDiagram: A ‘ggplot2’ Implement of Venn Diagram. https://CRAN.R-project.org/package=ggVennDiagram

Ghosh, R., Bloom, J. S., Mohammadi, A., Schumer, M. E., Andolfatto, P., Ryu, W., & Kruglyak, L. (2015, Aug). Genetics of Intraspecies Variation in Avoidance Behavior Induced by a Thermal Stimulus in Caenorhabditis elegans. Genetics, 200(4), 1327–1339. https://doi.org/10.1534/genetics.115.178491

Grishok, A., Pasquinelli, A. E., Conte, D., Li, N., Parrish, S., Ha, I., Baillie, D. L., Fire, A., Ruvkun, G., & Mello, C. C. (2001, Jul 13). Genes and mechanisms related to RNA interference regulate expression of the small temporal RNAs that control C. elegans developmental timing. Cell, 106(1), 23–34. https://doi.org/10.1016/s0092-8674(01)00431-7

Hahnel, S. R., Zdraljevic, S., Rodriguez, B. C., Zhao, Y., McGrath, P. T., & Andersen, E. C. (2018, Oct). Extreme allelic heterogeneity at a Caenorhabditis elegans beta-tubulin locus explains natural resistance to benzimidazoles. PLoS Pathog, 14(10), e1007226. https://doi.org/10.1371/journal.ppat.1007226

Harris, T. W., Arnaboldi, V., Cain, S., Chan, J., Chen, W. J., Cho, J., Davis, P., Gao, S., Grove, C. A., Kishore, R., Lee, R. Y. N., Muller, H. M., Nakamura, C., Nuin, P., Paulini, M., Raciti, D., Rodgers, F. H., Russell, M., Schindelman, G., Auken, K. V., Wang, Q., Williams, G., Wright, A. J., Yook, K., Howe, K. L., Schedl, T., Stein, L., & Sternberg, P.W. (2020, Jan 8). WormBase: a modern Model Organism Information Resource. Nucleic Acids Res, 48(D1), D762–D767. https://doi.org/10.1093/nar/gkz920

He, F. (2011, 2011/03/20). Total RNA Extraction from C. elegans. Bio-protocol, 1(6), e47. https://doi.org/10.21769/BioProtoc.47

Kamath, R. S., & Ahringer, J. (2003, Aug). Genome-wide RNAi screening in Caenorhabditis elegans. Methods, 30(4), 313–321. https://doi.org/10.1016/s1046-2023(03)00050-1

Kamath, R. S., Martinez-Campos, M., Zipperlen, P., Fraser, A. G., & Ahringer, J. (2001). Effectiveness of specific RNA-mediated interference through ingested double-stranded RNA in Caenorhabditis elegans. Genome Biol, 2(1), RESEARCH0002. https://doi.org/10.1186/gb-2000-2-1-research0002

Larsson, J. (2021). eulerr: Area-Proportional Euler and Venn Diagrams with Ellipses. https://CRAN.R-project.org/package=eulerr

Lee, D., Zdraljevic, S., Stevens, L., Wang, Y., Tanny, R. E., Crombie, T. A., Cook, D. E., Webster, A. K., Chirakar, R., Baugh, L. R., Sterken, M. G., Braendle, C., Felix, M. A., Rockman, M. V., & Andersen, E. C. (2021, Jun). Balancing selection maintains hyper-divergent haplotypes in Caenorhabditis elegans. Nat Ecol Evol, 5(6), 794–807. https://doi.org/10.1038/s41559-021-01435-x

Lee, S. H., Wong, R. R., Chin, C. Y., Lim, T. Y., Eng, S. A., Kong, C., Ijap, N. A., Lau, M. S., Lim, M. P., Gan, Y. H., He, F. L., Tan, M. W., & Nathan, S. (2013, Sep 10). Burkholderia pseudomallei suppresses Caenorhabditis elegans immunity by specific degradation of a GATA transcription factor. Proc Natl Acad Sci U S A, 110(37), 15067–15072. https://doi.org/10.1073/pnas.1311725110

Li, H.-D. (2018). GTFtools: a Python package for analyzing various modes of gene models. bioRxiv, 263517. https://doi.org/10.1101/263517

Love, M. I., Huber, W., & Anders, S. (2014). Moderated estimation of fold change and dispersion for RNA-seq data with DESeq2. Genome Biol, 15(12), 550. https://doi.org/10.1186/s13059-014-0550-8

McGrath, P. T., Rockman, M. V., Zimmer, M., Jang, H., Macosko, E. Z., Kruglyak, L., & Bargmann, C. I. (2009, Mar 12). Quantitative mapping of a digenic behavioral trait implicates globin variation in C. elegans sensory behaviors. Neuron, 61(5), 692–699. https://doi.org/10.1016/j.neuron.2009.02.012

Na, H., Zdraljevic, S., Tanny, R. E., Walhout, A. J. M., & Andersen, E. C. (2020, Aug). Natural variation in a glucuronosyltransferase modulates propionate sensitivity in a C. elegans propionic acidemia model. PLoS Genet, 16(8), e1008984. https://doi.org/10.1371/journal.pgen.1008984

Neuwirth, E. (2022). RColorBrewer: ColorBrewer Palettes. https://CRAN.R-project.org/package=RColorBrewer

Paaby, A. B., White, A. G., Riccardi, D. D., Gunsalus, K. C., Piano, F., & Rockman, M. V. (2015, Aug 22). Wild worm embryogenesis harbors ubiquitous polygenic modifier variation. Elife, 4. https://doi.org/10.7554/eLife.09178

Patro, R., Duggal, G., Love, M. I., Irizarry, R. A., & Kingsford, C. (2017, Apr). Salmon provides fast and bias-aware quantification of transcript expression. Nat Methods, 14(4), 417–419. https://doi.org/10.1038/nmeth.4197

Pedersen, B. S., & Quinlan, A. R. (2018, Mar 1). Mosdepth: quick coverage calculation for genomes and exomes. Bioinformatics, 34(5), 867–868. https://doi.org/10.1093/bioinformatics/btx699

Pertea, G., & Pertea, M. (2020). GFF Utilities: GffRead and GffCompare. F1000Res, 9. https://doi.org/10.12688/f1000research.23297.2

Pollard, D. A., & Rockman, M. V. (2013, Jun 21). Resistance to germline RNA interference in a Caenorhabditis elegans wild isolate exhibits complexity and nonadditivity. G3 (Bethesda, Md.), 3(6), 941-947. https://doi.org/10.1534/g3.113.005785

R Core Team. (2021). R: A language and environment for statistical computing. R Foundation for Statistical Computing. https://www.R-project.org/

Rockman, M. V., Skrovanek, S. S., & Kruglyak, L. (2010, Oct 15). Selection at linked sites shapes heritable phenotypic variation in C. elegans. Science, 330(6002), 372–376. https://doi.org/10.1126/science.1194208

Saber, S., Snyder, M., Rajaei, M., & Baer, C. F. (2022, May 6). Mutation, selection, and the prevalence of the Caenorhabditis elegans heat-sensitive mortal germline phenotype. G3 (Bethesda, Md.), 12(5). https://doi.org/10.1093/g3journal/jkac063

Sijen, T., Fleenor, J., Simmer, F., Thijssen, K. L., Parrish, S., Timmons, L., Plasterk, R. H., & Fire, A. (2001, Nov 16). On the role of RNA amplification in dsRNA-triggered gene silencing. Cell, 107(4), 465–476. https://doi.org/10.1016/s0092-8674(01)00576-1

Soneson, C., Love, M. I., & Robinson, M. D. (2015). Differential analyses for RNA-seq: transcript-level estimates improve gene-level inferences. F1000Res, 4, 1521. https://doi.org/10.12688/f1000research.7563.2

Stephens, M. (2017, Apr 1). False discovery rates: a new deal. Biostatistics, 18(2), 275–294. https://doi.org/10.1093/biostatistics/kxw041

Stiernagle, T. (2006, Feb 11). Maintenance of C. elegans. WormBook, 1-11. https://doi.org/10.1895/wormbook.1.101.1

Tijsterman, M., Okihara, K. L., Thijssen, K., & Plasterk, R. H. (2002, Sep 3). PPW-1, a PAZ/PIWI protein required for efficient germline RNAi, is defective in a natural isolate of C. elegans. Curr Biol, 12(17), 1535–1540. https://doi.org/10.1016/s0960-9822(02)01110-7

Vinuela, A., Snoek, L. B., Riksen, J. A., & Kammenga, J. E. (2010, Jul). Genome-wide gene expression regulation as a function of genotype and age in C. elegans. Genome Res, 20(7), 929–937. https://doi.org/10.1101/gr.102160.109

Webster, A. K., Hung, A., Moore, B. T., Guzman, R., Jordan, J. M., Kaplan, R. E. W., Hibshman, J. D., Tanny, R. E., Cook, D. E., Andersen, E., & Baugh, L. R. (2019). Population Selection and Sequencing of Caenorhabditis elegans Wild Isolates Identifies a Region on Chromosome III Affecting Starvation Resistance. G3 (Bethesda, Md.), 9(10), 3477-3488. Retrieved 2019/10//, from http://europepmc.org/abstract/MED/31444297 https://doi.org/10.1534/g3.119.400617 https://europepmc.org/articles/PMC6778785 https://europepmc.org/articles/PMC6778785?pdf=render

Wickham, H. (2016). ggplot2: Elegant Graphics for Data Analysis. Springer-Verlag New York. https://ggplot2.tidyverse.org

Wilke, C. O. (2020). cowplot: Streamlined Plot Theme and Plot Annotations for ‘ggplot2’. https://CRAN.R-project.org/package=cowplot

Wilson, R. C., & Doudna, J. A. (2013). Molecular mechanisms of RNA interference. Annu Rev Biophys, 42, 217–239. https://doi.org/10.1146/annurev-biophys-083012-130404

Zamanian, M., Cook, D. E., Zdraljevic, S., Brady, S. C., Lee, D., Lee, J., & Andersen, E. C. (2018, Mar). Discovery of genomic intervals that underlie nematode responses to benzimidazoles. PLoS Negl Trop Dis, 12(3), e0006368. https://doi.org/10.1371/journal.pntd.0006368

Zan, Y., Shen, X., Forsberg, S. K., & Carlborg, O. (2016, Aug 9). Genetic Regulation of Transcriptional Variation in Natural Arabidopsis thaliana Accessions. G3 (Bethesda, Md.), 6(8), 2319-2328. https://doi.org/10.1534/g3.116.030874

Zdraljevic, S., Fox, B. W., Strand, C., Panda, O., Tenjo, F. J., Brady, S. C., Crombie, T. A., Doench, J. G., Schroeder, F. C., & Andersen, E. C. (2019, Apr 8). Natural variation in C. elegans arsenic toxicity is explained by differences in branched chain amino acid metabolism. Elife, 8. https://doi.org/10.7554/eLife.40260

Zdraljevic, S., Strand, C., Seidel, H. S., Cook, D. E., Doench, J. G., & Andersen, E. C. (2017, Jul). Natural variation in a single amino acid substitution underlies physiological responses to topoisomerase II poisons. PLoS Genet, 13(7), e1006891. https://doi.org/10.1371/journal.pgen.1006891

Zhang, G., Roberto, N. M., Lee, D., Hahnel, S. R., & Andersen, E. C. (2022, Jun 16). The impact of species-wide gene expression variation on Caenorhabditis elegans complex traits. Nat Commun, 13(1), 3462. https://doi.org/10.1038/s41467-022-31208-4

